# Topoisomerase I is an Evolutionarily Conserved Key Regulator for Satellite DNA Transcription

**DOI:** 10.1101/2024.05.03.592391

**Authors:** Zhen Teng, Lu Yang, Qian Zhang, Yujue Chen, Xianfeng Wang, Yiran Zheng, Aiguo Tian, Di Tian, Zhen Lin, Wu-Min Deng, Hong Liu

## Abstract

Repetitive satellite DNAs, divergent in nucleic-acid sequence and size across eukaryotes, provide a physical site for centromere assembly to orchestrate chromosome segregation during the cell cycle. These non-coding DNAs are transcribed by RNA polymerase (RNAP) II and the transcription has been shown to play a role in chromosome segregation, but a little is known about the regulation of centromeric transcription, especially in higher organisms with tandemly-repeated-DNA-sequence centromeres. Using RNA interference knockdown, chemical inhibition and AID/IAA degradation, we show that Topoisomerase I (TopI), not TopII, promotes the transcription of α-satellite DNAs, the main type of satellite on centromeres in human cells. Mechanistically, TopI localizes to centromeres, binds RNAP II and facilitates RNAP II elongation on centromeres. Interestingly, in response to DNA double-stranded breaks (DSBs) induced by chemotherapy drugs or CRSPR/Cas9, α-satellite transcription is dramatically stimulated in a DNA damage checkpoint-independent but TopI-dependent manner. These DSB-induced α-satellite RNAs were predominantly derived from the α-satellite high-order repeats of human centromeres and forms into strong speckles in the nucleus. Remarkably, TopI-dependent satellite transcription also exists in mouse 3T3 and *Drosophila* S2 cells and in *Drosophila* larval imaginal wing discs and tumor tissues. Altogether, our findings herein reveal an evolutionally conserved mechanism with TopI as a key player for the regulation of satellite transcription at both cellular and animal levels.

## Introduction

Chromosome segregation during the cell division is essential for imparting the genetic information stored in chromosomes from one generation to another. Errors in this process will not only impair the normal development and growth of organisms, but also result in many diseases, such as cancer and infertility^1^. Chromosome segregation is orchestrated by the evolutionally conserved centromere^2^, which is usually built upon satellite DNAs for most eukaryotes. Satellite DNAs, one of the most divergent non-coding regions on a chromatin across eukaryotes^3^, had once been thought to be transcriptionally inert but are now known to undergo active transcription catalyzed by RNA Polymerase (RNAP) II^4,5^. The RNAP II-dependent satellite transcription promotes centromere function by maintaining centromere identity across eukaryotes^6–13^ and centromeric cohesion in human cells^14–17^. Both ongoing transcriptional process per se and transcribed non-coding satellite RNAs seem to be involved. However, the intrinsic and external responsive regulations of satellite transcription are little understood, especially in higher organisms including human, in which the centromere typically forms at hundreds to thousands of α-satellite DNA repeats^18^, which are further assembled into high-order repeats (HORs). This could be partially due to lack of accurate and complete centromere sequencing and assembly^17^. Fortunately, the most recent advance in complete human genome sequencing and assembly offers a grant opportunity for us to better understand the role and regulation of α-satellite transcription^19–21^. By taking the advantage of this new genome sequence information and performing a set of cellular, biochemical, and animal studies using human, mouse and *Drosophila* model systems, we intended to determine whether the regulatory mechanisms for RNAP II transcription on such evolutionarily divergent satellite DNAs are conserved and what might be the key players in this under-studied transcription of satellite repeats. Our findings herein reveal an evolutionally conserved mechanism with Topoisomerase I (TopI) as a key player for the regulation of satellite transcription at both cellular and animal levels.

## Results

### TopI is required for α-satellite transcription in human cells

In order to understand the regulation of α-satellite transcription, we concentrated on the protein factors that have been demonstrated to play dual roles in transcription and centromere functions. Topoisomerases are the enzymes that regulate transcription by managing DNA supercoils ^22^ and are enriched (TopI and II) on centromeres^23–25^. After treating Log-phase HeLa cells with TopI inhibitors camptothecin (CPT) and topotecan (TPT), we extracted total RNAs for real-time PCR analysis using two pairs of primers for housekeeping genes, GAPDH and RPL30, and three pairs of representative centromere primers, α-satellite1 (α-Sat1), α-satellite4 (α-Sat4) and α-satellite13/21 (α-Sat13/21), as previously described^15^. These primers amplify different but similar types of α-satellite RNAs. By comparing Ct values, we found that the expression levels of α-Sat1 (Ct, ∼30), α-Sat4 (Ct, ∼24) and α-Sat13/21 (Ct, ∼20) RNAs are from low and medium to high. In comparison, the levels of α-Sat13/21 RNA are much lower than the levels of housekeeping GAPDH mRNA (Ct, 12) but are still comparable to the levels of RPL30 mRNAs (Ct, ∼20), a ribosomal subunit. However, given high copy number of α-satellite DNAs, the expression levels of α-SatRNAs are therefore considered low. Notably, α-SatRNAs analyzed by these primers may represent RNAs produced from a broader region on α-satellite high-order repeats (HORs) than the CENP-A region. We found that both CPT and TPT treatment for 12 hrs significantly decreased the levels of α-Sat4 and α-Sat13/21 RNAs without changing much the levels of GAPDH and RPL30 mRNAs (**Fig. 1a** and **Supplementary Fig. 1a**). These decreases started at 6 hrs after CPT treatment (**Figs. 1a** and **1b**). We also noticed that CPT treatment did not significantly decrease the expression levels of α-Sat1 RNA, a lowly expressed α-satellite. By examining more regions of centromere 1, we found that the transcription on distinct regions within the same centromere displayed differential sensitivity to TopI inhibition **(Fig. 1b**). RNAi interference experiments also demonstrated that TopI knockdown significantly decreased the levels of all tested α-SatRNAs (**Fig. 1c** and **Supplementary Fig. 1c**). To further confirm the role of TopI in α-satellite transcription, we knocked in mAID to TopI C-terminus in HeLa cells stably expressing adapter protein Tir1 using CRISPR/Cas9 ^26^ (**Supplementary Fig. 2a**). Addition of IAA dramatically decreased the levels of TopI-mAID (**Supplementary Fig. 2b** and **c**), validating the effectiveness of this system. Interestingly, the degradation of TopI-mAID also significantly decreased the levels of α-Sat4 and α-Sat13/21 RNAs (**Fig. 1d**). Thus, with three distinct assays of TopI inhibitors, siRNAs and AID-degradation, we have revealed a novel role of TopI in α-satellite transcription, at least in several representative types of α-satellite DNAs. As the centromere on each chromosome contains different types of α-satellite high-order repeats (HORs) that are assembled from monomeric α-satellite, we wanted to know to what extent TopI is required for α-satellite transcription globally. We therefore performed RNA-seq analysis to assess the transcription on all the recently annotated α-satellite HORs^19–21^. Remarkably, CPT treatment decreased the transcription for the majority of HORs with detected transcripts, ∼80% for HeLa cells and ∼95% for RPE-1 cells. On average, CPT treatment decreased the HOR transcription by ∼75% for both HeLa and RPE-1 cells (**Figs. 1e, 3f** and **3g**). Thus, These RNA-seq results not only validate the findings from our real-time PCR analyses, but also support the notion that TopI is a global key regulator for the transcription of α-satellite DNAs in human genome. As α-satellite is being transcribed at a low level, we wanted to know whether Top I is also responsible for the transcription of lowly-expressed genes. By analyzing a group of lowly-expressed genes, whose TPM each had less than 2 TPM (Transcripts Per Million) as opposed to an average of 16 TPM per gene globally, we found that CPT treatment significantly affected the expression of only ∼10% of lowly expressed genes (**Supplementary Fig. 9c**), suggesting that TopI may not be a factor generally required for the expression of lowly transcribed genes. We next sought to determine whether TopI is also required for the transcription of other types of repetitive DNA sequences. Surprisingly, CPT treatment barely affected the transcription of ribosomal repeats and increased the transcription of telomeric repeats (**Fig. 1f**), highlighting the uniqueness of TopI in the transcription of α-satellite repeats. Moreover, by comparing RNAP II inhibitor α-amanitin and TopI inhibitor CPT, we found that CPT treatment efficiently repressed α-satellite transcription although α-amanitin seemingly exhibited a slightly higher efficiency (**Fig. 1g**).

**Figure 1.**
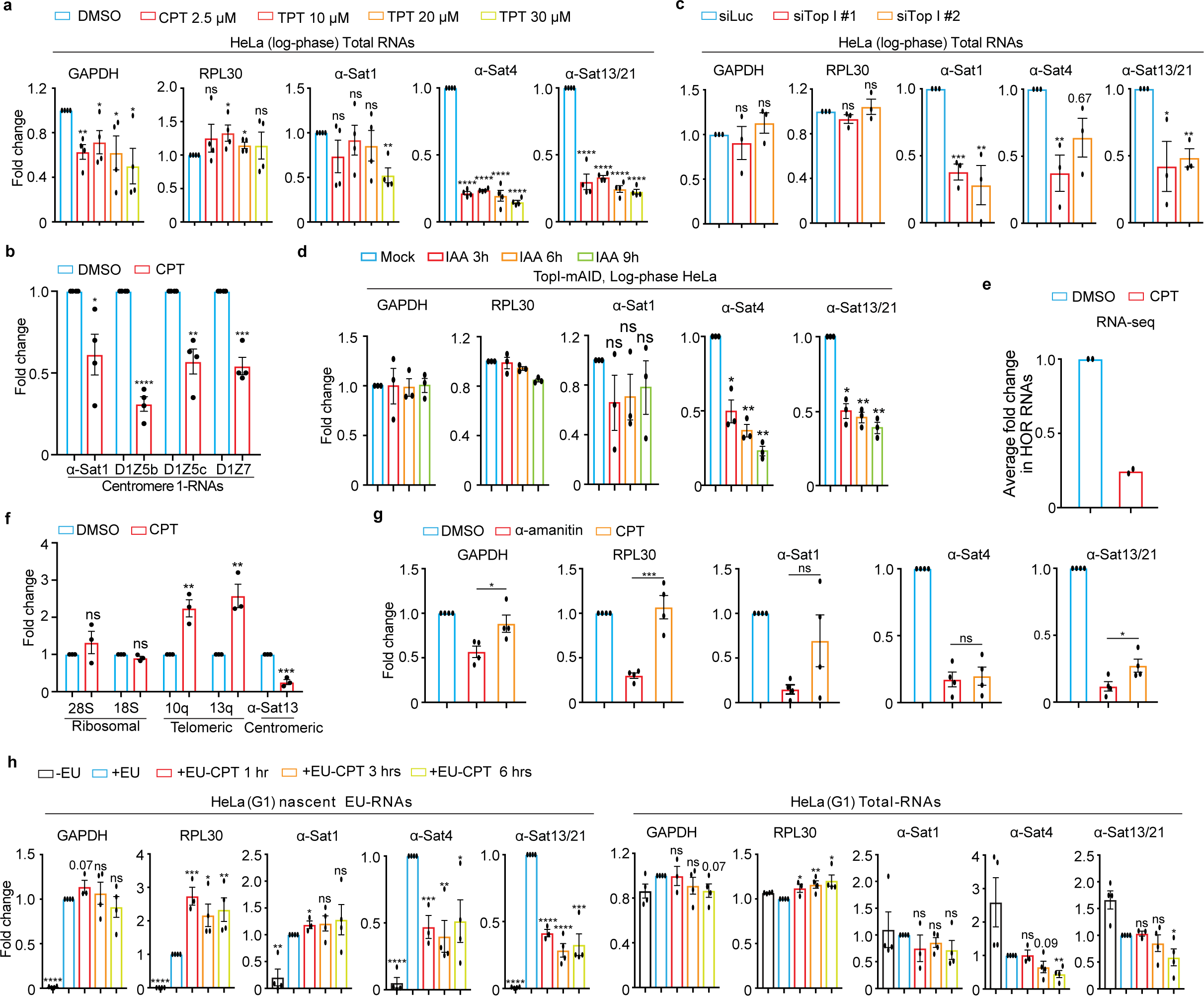
TopI is for α-satellite transcription in human cells. **A. TopI inhibition decreases the levels of α-SatRNAs.** Log-phase HeLa cells were treated with DMSO, CPT (2.5 µM) or TPT at distinct doses for 12 hrs and total RNAs were extracted for real-time PCR analysis with the indicated primers. **B. Effects of TopI inhibition on the transcription on different regions of centromere 1**. Log-phase HeLa cells were treated with DMSO or CPT (2.5 µM) and total RNAs were extracted for real-time PCR analysis. **C. TopI knockdown decreases the levels of α-SatRNAs.** HeLa cells were transfected with Luciferase or Top1 siRNAs (#1 and #2) and total RNAs were extracted for real-time PCR analysis. **D. IAA-induced TopI-mAID degradation decreases the levels of α-SatRNAs.** HeLa cells of TopI-mAID constructed by CRSPR/Cas9 were treated with IAA (1 µM) for the indicated time. Total RNAs were extracted for real-time PCR analysis. **E. TopI inhibition globally decreases the levels of α-satellite high-order repeat (HOR) RNAs.** HeLa cells treated with DMSO or CPT (2.5 µM) for 12 hrs and total RNAs were extracted for RNA-Seq analysis. Fold change for the transcripts of each HORs upon CPT treatment was calculated and the average of fold changes was shown here. **F. TopI inhibition does not decrease the transcription of telomeric and ribosomal repeats.** Log-phase HeLa cells were treated with DMSO or CPT (2.5 µM) for 12 hrs and RNAs were extracted for real-time PCR analysis. **G. Comparison in the effects of TopI inhibition and RNAP II inhibition on α-satellite transcription.** HeLa cells were treated with DMSO, α-amanitin (50 µg/ml) and CPT (2.5 µM) for 12 hrs, and RNAs were extracted for real-time PCR analysis. H**. TopI inhibition represses transcriptional activity on the centromere in G1 cells.** Thymidine-arrested G1 HeLa cells were treated with DMSO or CPT (2.5 µM) at different timepoints and 5′-ethynyl uridine (EU) was added 1 hr before harvest. Total RNAs and EU-RNAs were purified for real-time PCR analysis.

To rule out possible cell-cycle effects of TopI inhibitors and siRNAs on α-satellite transcription, we determined how α-satellite transcription is regulated during the cell cycle by analyzing α-SatRNAs in HeLa cells arrested with thymidine (G1), RO-3306 (G2) and nocodazole (M). Real-time PCR analyses demonstrated that α-satellite transcription is loosely cell-cycle regulated with a peak at G1 phase and a valley at M phase (**Supplementary Fig. 3a**). Flow-cytometric analyses demonstrated that CPT treatment for 12 hrs increased G1 population by ∼15% and decreased G2 population by ∼15% in HeLa cells and barely did so in RPE-1 cells (**Supplementary Fig. 3b** and **c**). TopI siRNA treatment for 48 hrs did not alter cell cycle profile either (**Supplementary Fig. 3e**). Thus, change in cell-cycle profile unlikely contributes significantly to decreased α-satellite transcription by TopI inhibition or knockdown. To further confirm this, we examined to what extent TopI is required for α-satellite transcription in thymidine-arrested G1 cells with higher centromeric transcription. Strikingly, CPT treatment for as short as 1 hr efficiently repressed transcriptional activity on α-satellite DNAs, as revealed by 5′-ethynyl uridine (EU)-labelled nascent RNAs, while total α-SatRNAs had not started decreasing until 6 hrs of etoposide treatment (**Fig. 1h** and **Supplementary Fig. 3d**). These results rule out cell-cycle effects of these treatments on α-satellite transcription, and further confirm a critical role of TopI in α-satellite transcription.

### Intact nucleoli and active ribosomal biogenesis is dispensable for TopI-mediated α-satellite transcription

We also examined to what extent TopI regulates α-satellite transcription through the nucleolus as TopI localizes to nucleolus and the nucleolus has been linked to α-satellite transcription ^22,27^. Using a potent RNAPI inhibitor BMH-21 ^28^, we found that the nucleoli in HeLa cells were dramatically fragmentated accompanied with a significant decrease in the levels of two major nucleolar component nucleolin and B23 (**Supplementary Fig. 4a** and **b**). Under this conduction, the levels of α-Sat4 and α-Sat13/21 RNAs slightly increased (**Supplementary Fig. 4c**), consistent with a repressive role of the nucleolus in α-satellite transcription proposed in a recent report ^27^. Remarkably, further CPT treatment for cells with fragmentated nucleoli still largely decreased α-satellite transcription, just as it did for cells with intact nucleoli (**Supplementary Fig. 4c**). Thus, these results suggest that the intact nucleolus and active ribosomal biogenesis may not play an important role in TopI-mediated α-satellite transcription, although we cannot rule out the possibility that some of nucleolar components may still regulate α-satellite transcription.

### TopI localizes to centromeres, physically interacts with RNAP II and is required for RNAP II elongation on α-satellite DNAs

Next, we asked how TopI promotes α-satellite transcription. Phosphorylation of RNAP II CTD at Serine2 (RNAP II-pSer2) is an important indicator for RNAP II elongation ^29^. We therefore examined the dependency of RNAP II elongation on TopI by conducting chromatin immunoprecipitation after treating log-phase HeLa cells with CPT. Compared to DMSO, CPT treatment dramatically decreased pSer2 levels by ∼80% on all the tested α-satellite regions, whereas it only slightly changed pSer2 levels on two housekeeping genes GAPDH and RPL30 (**Fig. 2a**). This result suggests that TopI is critical for active RNAP II elongation on α-satellite DNAs. To further confirm this, we performed florescence microscopy to examine how TopI inhibition affects pSer2 levels at specific cell cycle stages. RNAP II-pSer2 signals were robustly detected on the stretched centromeric chromatin in thymidine-arrested G1 HeLa cells (**Fig. 2b**) and were enriched on the centromere in nocodazole-arrested mitotic cells ^6,14,15,30,31^ (**Fig. 2c**). CPT treatment significantly decreased pSer2 levels on the centromere in both G1 and mitotic cells (**Figs. 2b** and **c**), which is consistent with our ChIP results (**Fig. 2a**). We next asked how TopI facilitates RNA II elongation. TopI was previously shown to physically interact with RNAP II to promote gene transcription ^32^. We then tested which isoform of RNAP II, phosphorylated or non-phosphorylated, was involved. Co-immunoprecipitation experiments verified the physical interaction between TopI and RNAP II in HeLa cells (**Fig. 2d, left panel**). Remarkably, the interaction appeared to occur only between TopI and phosphorylated RNAP II (pSer2) (**Fig. 2D, right panel)**, not between TopI and non-phosphorylated RNAP II ( **Supplementary Fig. 1d**), further supporting the notion that TopI binds RNAP II to facilitate RNAP II elongation on α-satellite DNAs. Notably, these physical interactions detected here were global, not restricted to α-satellite DNAs only. Finally, florescence microscopy demonstrated that TopI and its active intermediate TopI-cleavage complex were both present on the stretched centromeric chromatin in interphase cells (**Fig. 2e**). Strikingly, in mitosis, TopI-cc was enriched on the centromere without obvious detection on chromosome arms, whereas TopI was present along the entire chromosome including the centromere (**Figs. 2e** and **2f**). TopI and TopI-cc fluorescence signals were validated by siRNA knockdown (**Supplementary Fig. 1e** and **Sf**). TopI and TopI-cc localizing to centromeres provides further evidence to support the involvement of TopI in RNAP II transcription on α-satellite DNAs. Thus, we propose that TopI localizes to α-satellite DNAs and binds RNAP II to facilitate RNAP II elongation on α-satellite DNAs.

**Figure 2.**
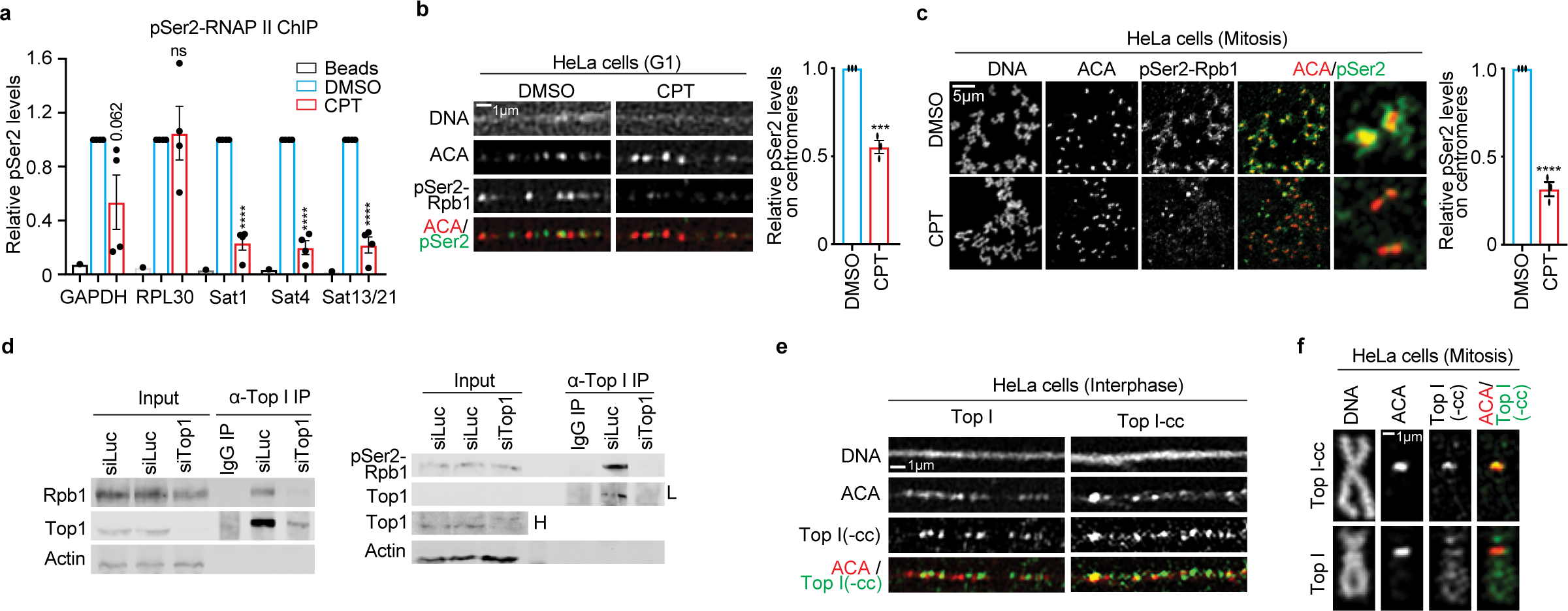
TopI binds RNAP II and promotes RNAP II elongation on α-satellite chromatin. **A. TopI inhibition decreases RNAP II-pSer2 levels on α-satellite DNAs.** HeLa cells treated with DMSO or CPT (2.5 µM) for 12 hrs were subjected to chromatin immunoprecipitation analysis with anti-RNAP II-pSer2 antibody. **B. TopI inhibition reduces RNAP II-pSer2 levels on α-satellite DNAs in G1 cells.** Stretched chromatin was prepared thymidine-arrested HeLa cells treated with DMSO or CPT (2.5 µM) for 6 hrs and immunostaining was performed. **C. Mitosis-specific inhibition of TopI reduces RNAP II-pSer2 levels on α-satellite DNAs.** Mitotic HeLa cells were enriched by a brief treatment of nocodazole (5 µM). Collected mitotic cells were further treated with DMSO or CPT (2.5 µM) for 5 hrs and then subjected for chromosome spread followed by immunostaining. **D. TopI physically interacts with RNAP II and phospho-RNAP II (pSer2).** Lysates of HeLa cells transfected Luciferase or Top1 siRNAs (#1) were incubated with IgG or antibody against TopI. Pelleted proteins were blotted with the indicated antibodies. The physical interaction between TopI and non-phospho-RNAP II was recorded in extended data fig.1e. “L” indicates low exposure and “H” high exposure. **E. TopI and TopI-cleavage complex (cc) are present on the centromere in interphase.** Stretched chromatin was prepared from log-phase HeLa cells and immunostaining was performed. **F. TopI-cc is enriched on the centromere in mitosis.** HeLa cells were briefly treated with nocodazole (5 µM) and mitotic cells were collected for chromosome spread followed by immunostaining. Validation of TopI and Top-cc fluorescence signals were recorded in Extended Data Fig. 1e,f. Average and standard error were usually shown in all the figures (here and hereafter) unless specified. Details of quantification and technical details in all the figures were recorded in the section of Methods and Materials. ns, not significant; *, P<0.05; **, P<0.01; ***, P<0.001; ****, P<0.0001.

### DNA double-stranded breaks (DSBs) dramatically stimulate α-satellite transcription in a DNA damage checkpoint-independent manner

We next determined whether TopII similarly regulates α-satellite transcription as TopI. We applied TopII inhibitor etoposide to HeLa cells for 12 hrs and then performed real-time PCR analysis. Unexpectedly, etoposide treatment increased the expression of the tested α-satRNAs, whereas it only had marginal impact on housekeeping GAPDH and RPL30 mRNAs (**Fig. 3a, left panel**). Such etoposide-induced α-satellite transcription seemed to be global, as revealed by real-time PCR analyses using primers against various types of α-satellite HORs from almost every centromere in HeLa cells (**Supplementary Fig. 5**). It was also noticed that the effects of etoposide on α-satRNA production varied a lot, ranging from several-fold to hundred-fold increase. By chasing cells with EU for 1 hr, we also found that etoposide treatment also dramatically increased the levels of nascent α-SatRNAs (**Fig. 3A, right panel)**. Thus, it is likely that etoposide treatment largely increases transcriptional activity on α-satellite, but we cannot completely rule out the possibility that altered dynamics of α-SatRNA stability by etoposide treatment also contributes to its dramatic increase. As etoposide not only inhibits TopII but also induces DNA double-stranded breaks (DSBs), we therefore asked which, TopII inhibition or DSBs, stimulated the production of α-SatRNAs using the following chemicals: ICRF (TopII inhibitor and no DSBs), teniposide (etoposide analog, TopII inhibitor and DSBs), phleomycin (non-TopII inhibitor and DSBs, and mitomycin C (MMC) and cisplatin (non-TopII inhibitor and no DSBs). Etoposide, phleomycin and MMC treatment for 12 hrs dramatically increased the levels of a known DNA-damage marker γH2AX throughout the nucleus including centromeres (**Supplementary Figs. 6a and b**), but only etoposide- and phleomycin-induced damage was DSBs, as proved by comet assay (**Fig. 3c**). Accordingly, etoposide, its analog teniposide and phleomycin dramatically induced α-satellite transcription, whereas ICRF, MMC and cisplatin, did not (**Figs. 3a, b, d, e**, and **Supplementary Figs. 6c-g**). TopII (a, b) depletion by siRNAs barely increased γH2AX levels and did not affect α-satellite transcription either (**Supplementary Figs 7a-c**). Similar results were also observed in non-transformed human RPE-1 cells (**Supplementary Figs. 8a and b**). Importantly, DSBs specifically generated by CPRSR/Cas9 on α-satellite DNAs were sufficient to increase α-satellite transcription (**Figs. 3f** and **g**). Thus, it is DSBs, but not TopII inhibition, that induce centromeric transcription. Notably, DSB-induced α-satellite transcription still largely depends on RNAP II, as revealed by two RNAP II inhibitors, α-amanitin and flavopiridol (**Supplementary Figs. 6h** and **i**).

**Figure 3.**
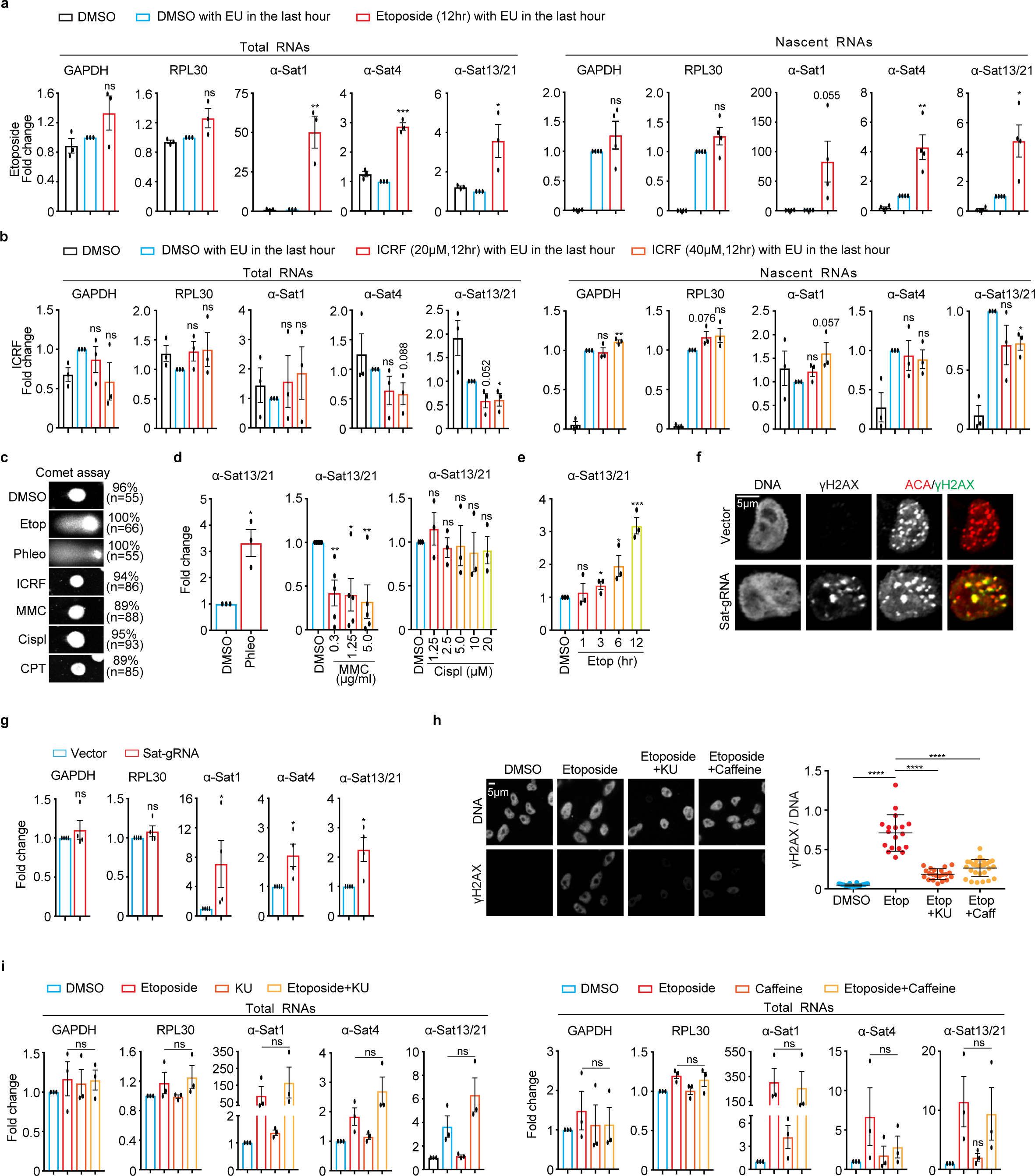
Double-stranded breaks (DSBs) dramatically increase α-satellite transcription in a DNA damage checkpoint-independent manner in human cells. **A** and **B. Treatment of etoposide (A), not ICRF (B), dramatically increases the levels of total and nascent α-satRNAs.** Total and EU-labelled RNAs extracted from log-phase HeLa cells treated with DMSO, etoposide (30 μM), or ICRF (20 or 40 μM), were subjected to real-time PCR analysis. **C. DSB analysis by comet assay.** HeLa cells treated with DMSO, Etoposide (Etop, 30 μM), Phleomycin (Phleo, 80 μg/ml), ICRF ( 20 μM), MMC (5 μg/ml), Cisplatin (Cispl, 20 μg/ml), or CPT (2.5 μM) for 12 hrs, were subjected to analysis by comet assay. **D. DSB-inducing agents increase α-satellite transcription.** HeLa cells were treated with DMSO, Phleo (80 μg/ml, 24 hrs), MMC (indicated, 12 hrs), or Cispl (indicated, 12 hrs) and RNAs were extracted for real-time PCR analysis. Result for α-Sat13/21 primer was shown here. Results for other primers were recorded in Figures S6D-6F. **E. Etoposide increases α-satellite transcription in a time-dependent manner.** HeLa cells were treated with DMSO or etoposide (2.5 μM) at the different timepoints and RNAs were extracted for real-time PCR analysis. Results for α-Sat13/21 primer were recorded here and results for GAPDH, RPL30 and α-Sat1 primers were recorded in Figure S6G. **F. CRISPR-Cas9 generates DSBs specifically on the centromere.** Plasmids containing Sat-gRNA were transfected into HeLa cells for 24 hrs and immunostaining was performed. ∼16% of cells had obvious γH2AX foci on centromeres at 24 hrs after expression of Sat-gRNA. **G. DSBs generated by CRISPR-Cas9 increase α-satellite transcription.** RNAs extracted from cells in (**E**) were subjected to real-time PCR analysis. **H. α-SatRNAs localize within the nucleus and their fluorescence signals increase over time of etoposide treatment.** HeLa cells treated with etoposide at the different timepoints were subjected to RNA-FISH analysis. Quantification of FISH signals (RNA-FISH/DNA) was shown in the bottom panel. **H. ATM inhibitor KU-55933 (KU) and caffeine treatment decrease γH2AX levels in etoposide-treated cells.** HeLa cells were treated with DMSO, etoposide (30 μM), etoposide plus KU-55933 (10 μM), or etoposide plus caffeine (2 μM) for 12 hrs and then stained with the indicated antibodies. Quantification of γH2AX levels was shown in the bottom panel. **I. ATM inhibitor KU or Caffeine treatment does not affect etoposide-induced α-satellite transcription.** RNAs extracted from cells in (**H**) were subjected to real-time PCR analysis. Average and standard deviation in (**H**, **right panel**) were shown.

We next tested whether DSB-induced α-satellite transcription is dependent on the DNA damage checkpoint using caffeine that can override the activation of the DNA damage checkpoint by inhibiting two key checkpoint kinases ATM and ATR ^33^. As expected, Caffeine treatment significantly reduced the etoposide-elevated γH2AX levels (**Fig 3h**), but it did not affect α-satellite transcription (**Fig 3i**, **right panel**). Nor was α-satellite transcription affected by ATM inhibition (KU-55933) (**Fig 3i, left panel**) or DNA-PK inhibition (NU7026) (**Supplementary Fig. 8e**). Moreover, DSB-induced α-satellite transcription seemed to be independent of the MRN complex (**Supplementary Fig. 8f**), which was shown to regulate DSB-associated RNA transcription^34^. Thus, DSB-induced α-satellite transcription is unlikely dependent on the DNA damage checkpoint.

### DSB-induced α-satellite transcription depends on TopI

We next asked whether DSB-induced α-satellite transcription is dependent on TopI since we have established TopI as a key regulator for α-satellite transcription in unperturbed cells. Consistently, etoposide treatment dramatically increased the levels of all tested α-SatRNAs in HeLa cells (**Fig. 4a**). Remarkably, simultaneous CPT treatment almost completely abolished the increase of α-SatRNAs induced by etoposide. Partial decrease of etoposide-induced α-SatRNAs were also observed in TopI-depleted HeLa cells (**Supplementary Fig. 8d**). In addition, similar results were observed in non-transformed human RPE-1 cells (**Fig 4b** and **Supplementary Fig. 8c**). Notably, treatment with these chemicals slightly changed the transcription of tested genes, but to a much less extent. These changes were unlikely caused by alteration in cell cycle profile, as our flow-cytometric analyses demonstrated that treatment of CPT or Etoposide only slightly changed it (**Supplementary Fig. 3c**). Furthermore, the degradation of TopI-mAID by IAA treatment also significantly decreased the levels of etoposide-induced α-SatRNAs (**Fig. 4d** and **Supplementary Fig. 2d**). Again, with three distinct approaches, we have provided strong evidence to support a key role of TopI in DSB-induced α-satellite transcription. By measuring EU-labelled nascent RNAs, we found that TopI-mediated changes of α-SatRNAs in both unperturbed and DNA-damaged cells were achieved through transcriptional activity (**Fig. 4c**). In addition, etoposide treatment barely affected the transcription of ribosomal repeats and increased the transcription of telomeric repeats, but the increase was independent of TopI (**Fig. 4e**). Taken these findings together, we conclude that TopI is a unique and key regulator for α-satellite transcription in response to DSBs.

**Figure 4.**
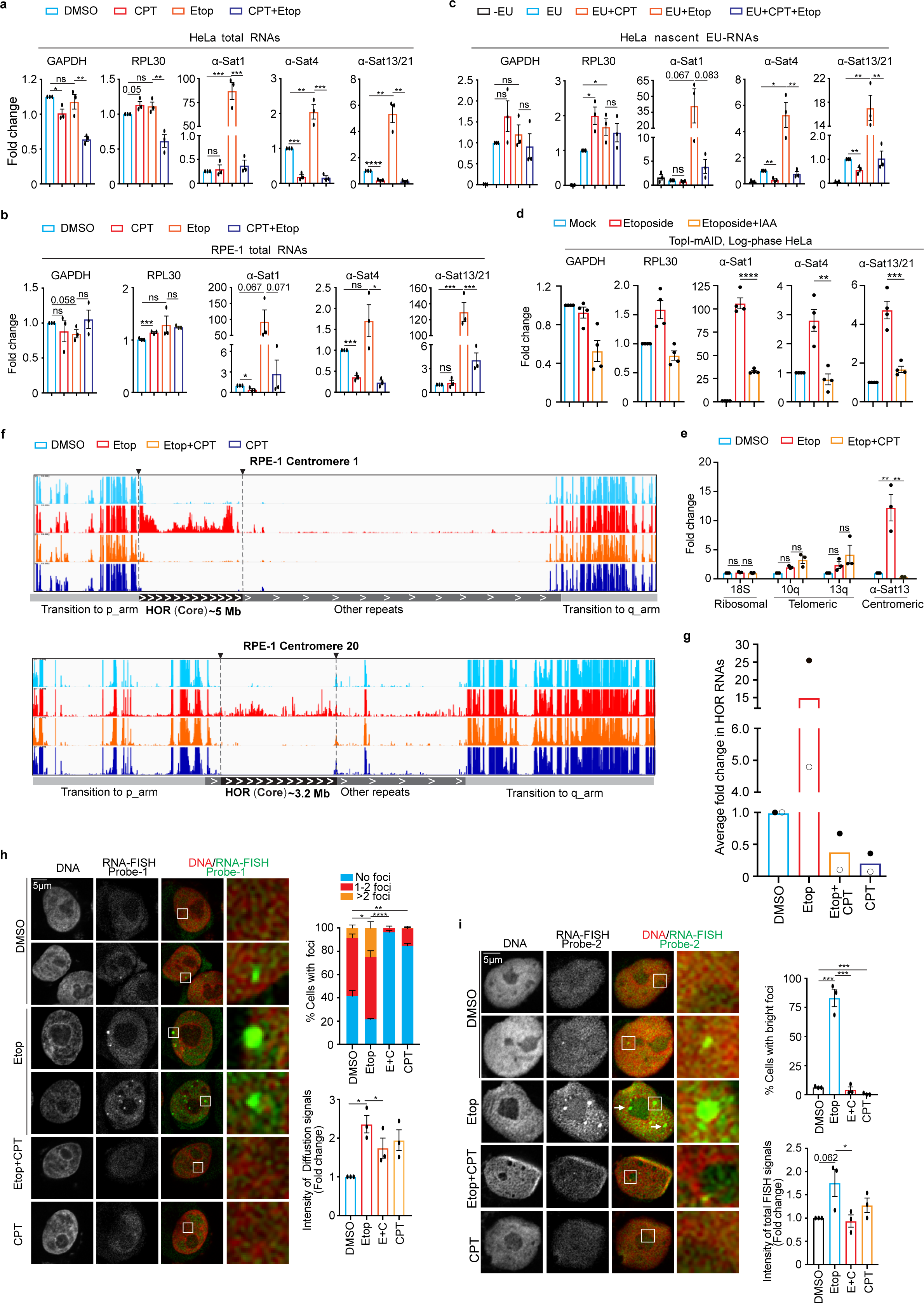
DSB-induced α-satellite transcription depends on TopI and increased RNAs are predominantly derived from α-satellite high-order-repeats (HORs) and spread across the nucleus in human cells. **A. TopI inhibition dramatically decreases the levels of total α-SatRNAs in etoposide-treated HeLa cells.** HeLa cells were treated with DMSO, CPT (2.5 μM), Etoposide (Etop, 30 μM), or CPT plus Etoposide for 12 hrs and total RNAs were extracted for real-time PCR analysis. **B. TopI inhibition dramatically reduces the levels of total α-SatRNAs in etoposide-treated RPE-1 cells.** RPE-1 cells were treated with DMSO, CPT (2.5 μM), Etoposide (30 μM), or Etoposide plus CPT (E+C) for 12 hrs, and total RNAs were extracted for real-time PCR analysis. **C. TopI inhibition abates the transcriptional activity in both unperturbed and etoposide-treated HeLa cells.** HeLa cells were treated with DMSO, CPT (2.5 μM), Etoposide (Etop, 30 μM), or Etoposide plus CPT for 12 hrs and EU was added 1 hr before harvest. EU-RNAs were purified for real-time PCR analysis. **D. IAA-induced TopI-mAID degradation decreases the levels of etoposide-induced α-SatRNAs.** HeLa cells of TopI-mAID were treated with IAA (1 μM) and etoposide for 9 hrs for the indicated time. Total RNAs were extracted for real-time PCR analysis with the indicated primers. **E. TopI is not required for the transcription of ribosomal repeats and telomeric repeats in etoposide-treated cells.** HeLa cells treated with DMSO, Etoposide (Etop, 30 μM), or Etoposide plus CPT (2.5 μM) for 12 hrs, and total RNAs were extracted for real-time PCR analysis. **F. RNA-sequencing (RNA-seq) analyses of α-satRNAs.** RPE-1 cells were treated with DMSO, CPT (2.5 μM), Etoposide (Etop, 30 μM), or CPT plus etoposide for 12 hrs and total RNAs were extracted for RNA-Seq analysis. Technical details were recorded in the section of Methods and Materials. **G. TopI inhibition globally decreases the levels of α-satellite high-order repeat (HOR) RNAs.** Fold change for the transcripts of each HOR transcript upon chemical treatment in (**E**) was calculated and the average of fold changes was shown. Average of two independent experiments (filled circular and non-filled circular dots) was shown here. **H** and **I. Fluorescence *in situ* hybridization (FISH) analysis of α-satRNAs.** HeLa cells were treated with DMSO, CPT (2.5 μM), Etoposide (Etop, 30 μM), or etoposide plus CPT for 12 hrs and then subjected to RNA-FISH analysis using probe-1 (**H**) and probe-2 (**I**). Quantifications of FISH foci signals (upper panels, **H** and **I**), diffusion signals (bottom panel, **H**) and total FISH signals (bottom, **I**) were shown here.

### DSB-increased α-SatRNAs are mainly derived from α-satellite high-order repeats (HORs)

To better understand how TopI inhibition globally affects α-satellite transcription, we performed RNA-seq by taking advantage of recently completed human centromere assembly ^19–21^. Each human centromere has a core region that contains evolutionarily youngest α-satellite high-order repeats (HORs) and decayed oldest α-satellite HORs ^20^. Within this core region resides the histone H3 variant CENP-A. Flanking to this core are the regions of monomeric α-satellite and other repetitive DNA sequences (hereafter, other repeats), and the transition regions containing expressed genes. After treating RPE-1 cells with DMSO, etoposide, CPT, or etoposide plus CPT for 12 hrs, we performed RNA-Seq analysis using a complete Telomere-to-Telomere reconstructed human reference genome ^21^. A relatively low-amplitude of transcription was detected at three representative centromeres 1, 10 and 20 (α-satellite HORs and other repeats), residing on a large-, medium- and small-sized chromosomes, respectively ^35^ (**Fig. 4F** and **Supplementary Fig. 9a** and **b**). Remarkably, etoposide treatment dramatically increased α-satRNA production and simultaneous treatment with CPT completely abrogated such increase (**Fig. 4g** and **Supplementary Fig. 9a**). Surprisingly, the etoposide-increased RNA production predominantly occurred throughout almost the entire α-satellite HORs (youngest and oldest), including but not limited to the CENP-A region, although slight changes were also observed on other repeats (**Figure 4f, Supplementary Figs. 5** and **9a**). This result is further confirmed by real-time PCR analysis using primers against different types of α-satellite HORs (**Supplementary Figs. 5**). CPT treatment alone also decreased the production of α-satellite HOR-derived RNAs (**Figs. 4f** and **4g**). In contrast to α-satellite transcription, these chemical treatments only significantly changed the expression of less than 2.1% of genes globally (**Methods and Materials**); nor did seemingly alter transcriptional profiles for three representative chromosomes 1, 10 and 20 (**Supplementary Figs. 9b**), suggesting that chemical treatments under our experimental conditions may not pose a significant change on global chromatin structure. Altogether, these findings further confirm that TopI is a key regulator for α-satellite transcription.

### DSB-increased α-SatRNAs mainly localize in the nucleus

α-SatRNAs have been found to localize at distinct places within cells: centromeres/kinetochores ^11,36–38^, nucleoli ^36,39^, and independent foci ^27^. We also sought to determine where DSB-increased centromere RNAs localize within the cell using RNA-FISH with fluorescein-labeled oligonucleotide probes. We then developed two RNA-FISH probes (probe-1 and probe-2). Our two probes detected two types of signals, foci and diffusion. Both types were largely diminished by RNase treatment (**Supplementary Figs. 10a**). We then used these probes to examine how DSBs and TopI inhibition affected the localization of α-satRNAs. Among DMSO-treated cells, ∼56% had 1-2 bright FISH foci for probe-1 and only ∼8% for probe-2, respectively (**Figs. 4h** and **i**). The difference may be due to distinct sensitivity of these two probes. Etoposide treatment significantly increased the intensity of diffusion signals throughout the nucleus by ∼2.2-fold for probe-1 and ∼1.7-fold for probe-2, respectively (**Figs. 4h** and **i**, **right panels**). Remarkably, accompanied by increased diffusion signals, the number of cells with foci signals and the intensity and number of foci within a cell were both significantly increased (**Figures 4h** and **i**). Further CPT-treatment almost abolished the etoposide-increased foci signals and moderately decreased the etoposide-elevated diffusion signals, which further validates the results from our real-time PCR and RNA-seq analyses. Thus, our results support the notion that α-SatRNAs mainly exist as foci in the nucleus but they can also appear as weak diffusion signals. Notably, FISH foci were not always present in the nucleolus (**Fig. 4h** and **i**), nor always co-localized with centromeres, as revealed by anti-centromere antibody (ACA) co-staining (**Supplementary Figs. 10b**).

### TopI-maintained satellite transcription is conserved in mouse and *Drosophila* cells

We next asked if the mechanism of TopI-maintained α-satellite transcription we have observed in human cells would also apply to other eukaryotic cells. In mouse 3T3 cells, CPT treatment for as short as 1 hr significantly inhibited the transcription of both minor and major satellites (**Fig. 5a**). Further siRNA knockdown of mouse TopI confirmed TopI is required for minor-satellite transcription (**Fig. 5b**). Consistently, etoposide treatment dramatically increased the amounts of centromere minor and major satellite RNAs by ∼80-fold and ∼25-fold, respectively, compared to DMSO treatment (**Fig. 5c**). These increases were significantly reduced by simultaneous CPT treatment. Thus, TopI-maintained satellite transcription is conserved in mouse cells. Similar results were also observed for satellite 359 (Sat359) RNA derived from the centromere of sex chromosome X In *Drosophila* S2 cells ^40^ (**Fig. 5d**), suggesting that TopI-maintained satellite transcription is also conserved in cultured *Drosophila* cells.

**Figure 5.**
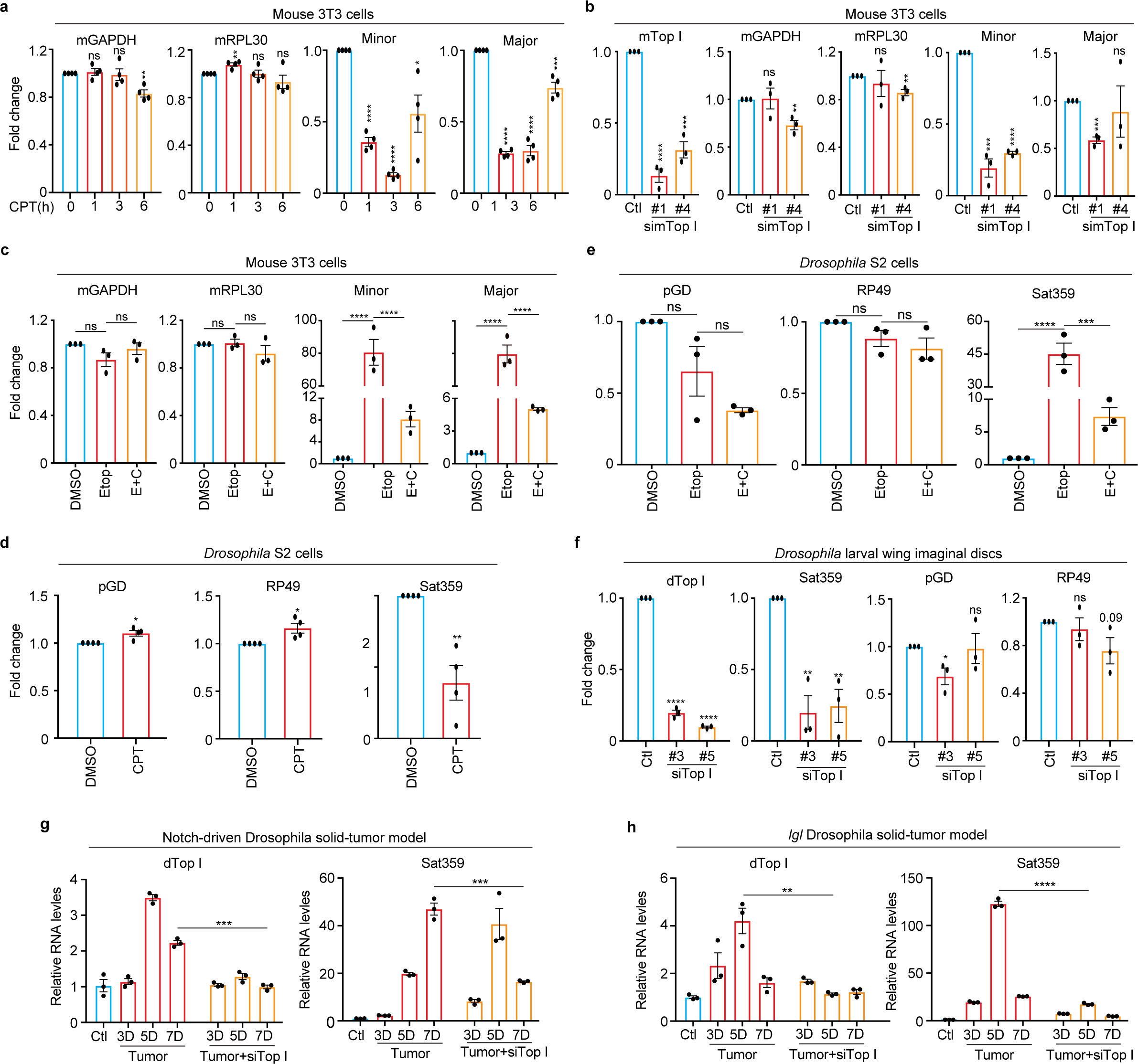
TopI-dependent satellite transcription is conserved across eukaryotes. **A. TopI inhibition decreases satellite transcription in mouse cells.** Mouse 3T3 cells were treated with CPT (2.5 μM) at the different timepoints, and RNAs were extracted for real-time PCR analysis. **B. TopI knockdown decreases satellite transcription in mouse cells.** Mouse 3T3 cells were transfected with mTopI siRNA (#1 and #2), and RNAs were extracted for real-time PCR analysis. **C. TopI is required for DSB-induced satellite transcription in mouse cells.** RNAs extracted from mouse 3T3 cells treated with DMSO, CPT (2.5 μM), Etoposide (Etop, 30 μM), or Etoposide plus CPT (E+C) for 12 hrs, were subjected to real-time analysis. **D. TopI inhibition decreases satellite transcription in *Drosophila* cells.** RNAs extracted from *Drosophila* S2 cells treated with DMSO or CPT (10 μM) for 6 or 12 hrs were subjected to real-time analysis. **E. TopI is required for DSB-induced satellite transcription in *Drosophila* cells.** RNAs extracted from *Drosophila* S2 cells treated with DMSO, CPT (10 μM), Etoposide (Etop, 30 μM), triptolide (Trip, 4 μM) or Etoposide plus CPT (E+C) for 12 hrs, were subjected to real-time analysis. **F. TopI promotes satellite transcription in *Drosophila* larvae tissues.** RNAs extracted from *Drosophila* larvae wing imaginal discs with mock or siTopI (#3 and #5) treatment were subjected to real-time PCR analysis. **G** and **H. Satellite transcription is gradually elevated in a TopI-dependent manner in *Drosophila* tumor tissues.** RNAs extracted from control tissues, notch-driven (**G**) or *lethal (s) giant larvae* (*lgl*) (**H**) *Drosophila* solid-tumor tissues with mock or siTopI treatment at different timepoints were subjected to real-time PCR analysis. The average and standard error here were calculated from three technical replicates.

### TopI-dependent satellite transcription is conserved in *Drosophila* larvae and tumor tissues

We next tested whether TopI-dependent satellite transcription is also conserved at the organismal level. Using an inducible *Act-Gal4/Gal80^ts^* system and TopI dsRNAs^41^, we knocked down TopI in *Drosophila* larvae and then assessed Sat359 RNA. Real-time PCR analyses on wing imaginal discs confirmed that TopI mRNA levels were largely reduced (**Fig. 5f**), accordingly, Sat359 RNA levels decreased by ∼70% without a notable impact on two housekeeping RP49 and pGD mRNAs. Thus, TopI-maintained satellite transcription is also a conserved mechanism in a developing tissue.

We then sought to determine whether TopI also regulates satellite transcription in cancer development. Notch-driven and *lethal (2) giant larvae (lgl) Drosophila* solid-tumor models were engaged as extensive DNA damage occurred in tumorigenesis and cancer development in these models can be monitored in a time-dependent manner^42–44^. Over time of tumor development, a trend of moderate increases (several-fold) in the expression of TopI mRNA was observed in both models (**Figs. 5g** and **h**); accordingly, satellite transcription also increased (up to a hundred-fold), even more dramatically. The dramatic increase of satellite transcription in these cancer tissues might be driven by extensive DNA damage. Remarkably, TopI knockdown by dsRNAs in these two models reduced the levels of Sat359 RNA (**Figs. 5g** and **h**). Thus, satellite transcription in cancer development is dependent, at least partially, on TopI in *Drosophila larvae*.

## Discussion

In budding yeast, satellite transcription seems to be repressed by centromere-binding factor cbf1 and histone H2A variant Htz1 ^45,46^. In *Drosophila*, FACT (facilitate chromatin transcription) has been shown to regulate the transcription on an ectopic centromere ^8^. A recent study suggested that centromere proteins may be involved in α-satellite transcription in human cells^27^. These factors are either species-specific or have never been demonstrated to conservatively regulate the transcription on divergent eukaryotic centromeres. It is therefore unknown whether universal regulators or mechanisms across eukaryotes exist to regulate satellite transcription. Using multiple approaches and different kinds of organisms, we have demonstrated that TopI is such a critical factor and TopI-mediated satellite transcription is a conserved mechanism. Specifically, RNA-seq analyses demonstrate that TopI is a global key regulator for α-satellite transcription in human cells. Mechanistically, TopI localizes to centromeres and physically interacts with RNAPII to promote RNAP II elongation on α-satellite DNAs. Such regulation seems to be more critical for α-satellite DNAs than for genes albeit the underlying mechanism is unknown. Remarkably, TopI’s impact on human α-satellite transcription is predominantly limited to the regions of α-satellite HORs in human centromeres. This is even more striking in response to DSBs. In the future, it would be of our interest to determine whether non-B type DNA structures formed on α-satellite might be one of the underlying mechanisms ^47^. It would also be intriguing to understand why TopI, not its relative TopII, is required for α-satellite transcription, although they both play a similar role in transcription. Notably, although TopI inhibition in our tests may not pose dramatic changes in chromatin structures, as revealed by our genome-wide transcriptional profiles, we cannot completely rule out the possibility that some minor changes in chromatin structures may also contribute to α-satellite transcription to a certain extent.

DSBs on genes usually slow down or block RNAP II transcription in an ATM- or DNA-PK-dependent manner, thus mitigating the threat to genomic stability imposed by transcription over DSBs ^48,49^. In contrast, we have here discovered a distinct mechanism, in which DSBs dramatically increase α-satellite transcription in an ATM- or DNA-PK-independent but a TopI-dependent manner. Interestingly, this novel mechanism is evolutionally conserved despite very divergent satellite DNA sequences across eukaryotes, further highlighting TopI as an evolutionally conserved key regulator for satellite transcription. Remarkably, we have also provided evidence to support that DSBs induce satellite transcription in cancer development in a TopI-dependent manner in Notch-driven and *lgl Drosophila* solid-tumor models, Thus, TopI-dependent satellite transcription exits at both cellular and animal levels. In addition, previous studies demonstrated that transcription on human satellite-III, not α-satellite, is induced in response to heat shock^50,51^, provoking an interesting question whether the transcription on distinct types of non-coding satellite DNA sequences is responsive to different types of stimuli. Although we have discovered a novel regulation of DSB-induced α-satellite transcription, the underlying mechanism is currently not understood. It will be of our great interest to address it in the future.

What is the function of satellite overexpression in response to DNA damage? Sat359 RNA in *Drosophila* was shown to be important for chromosome segregation^12^ and satellite RNA overexpression appeared to impair chromosomal stability in both human and murine cells^52^. It is therefore likely that DNA damage-increased satellite RNAs impair proper chromosome segregation during the cell division, thus affecting cell ploidy. In support of it, cells in the Notch-induced *Drosophila* solid-tumor model did suffer dramatic ploidy alterations in cancer development^42^. As aberrant overexpression of satellite repeats was found in human pancreatic and epithelial cancer tissues^53^, DNA damage-increased satellite transcription might play a role in tumorigenesis.

## Author Contributions

Conceptualization, H.L.; Methodology, Z.T., L.Y., Y.C., Q.Z., X. W., A.T., D.T., Z.L., W.D., and H.L.; Investigation, Z.T., L.Y., Q.Z., Y.C., X. W., Y.Z., A.T., Z.L.; Writing – Original Draft, H.L.; Writing – Review & Editing, Z.T., L.Y., Q.Z., Y.C., X. W., Y.Z., A.T., Z.L., W.D., and H.L.; Funding Acquisition, A.T., Z.L., W.D., and H.L.; Resources, A.T., W.D., and H.L.; Supervision, H.L..

## Data availability statement

Original RNA-seq data have been deposited to *NCBI SRA# SRP381962*. There are not any restrictions on data availability.

## Acknowledgment

This work was supported by Tulane University startup funds and the National Institute of General Medical Sciences/the National Institutes of Health grants R01GM124018 and R01GM141123, awarded to H. Liu. Research reported in this study was also supported by the National Cancer Institute of the National Institutes of Health under Award Number R01CA261258 to Z. Lin. The content is solely the responsibility of the authors and does not necessarily represent the official views of the National Institutes of Health.

The authors declare no competing financial interests

## Materials and Methods

### Cell culture, transfection, and chemicals

HeLa (Tet-On) cells purchased from Invitrogen were incubated in Dulbecco’s modified Eagle’s medium (DMEM, Invitrogen) containing 10% fetal bovine serum (FBS) and 10 mM L-glutamine at 37°C and in 5% CO_2_. Human RPE-1 cells (a gift from Dr. Hongtao Yu) were cultured in DMEM: F-12 medium (Invitrogen) supplemented with 10% FBS and 10mM L-glutamine at 37°C and in 5% CO_2_. Mouse NIH3T3 cells were raised with DMEM formulated with addition of 10% FBS at 37°C and in 5% CO_2_. Drosophila S2 cells were cultured in Schneider’s Medium (Invitrogen) at room temperature (23°C) supplemented with 10% FBS.

Chemicals used in this study: Nocodazole (M1404, Sigma Aldrich), α-amanitin (MilliporeSigma, A2263), Flavopiridol (Selleckchem, S1230), Triptolide (MilliporeSigma, T3652), Etoposide (MilliporeSigma, 341205), Camptothecin (CPT) (Cell signaling, 13637S), Topotecan (TPT) (MilliporeSigma, T2705), KU-55933 (MilliporeSigma, SML1109), Caffeine (MilliporeSigma, C0750), NU7026 (Selleckchem, S2893), Phleomycin (MilliporeSigma, P9564), Teniposide (MilliporeSigma, SML0609), ICRF (MilliporeSigma, I4695), Mitomycin C (APEXBio, A4452), Mitoxantrone (MTX, Sigma, M6545), Cisplatin (Simga, P4394), RO3306 (Sigma, SML0569), Thymidine (Sigma, T1895). These chemicals were dissolved in DMSO and working concentrations were as follows: Nocodazole, 5 μM; α-amanitin, 25 μg/ml; Flavopiridol, 1 μM; Etoposide, 30 μM; Camptothecin (CPT), 2.5 or 10 μM; Topotecan (TPT), 10-30 μM; KU-55933, 5-10 μM; caffeine, 1-2 uM; NU7206, 0.8 μM; Phleomycin, 80 μg/ml; Teniposide, 2.5 μM; ICRF, 20 μM; Mitomycin C (MMC), 5 μg/ml; Mitoxantrone (MTX), 2 μM; Cisplatin, 1.25-20 μM. Triptolide, 4 μM.

For RNAi experiments, siRNA oligonucleotides were purchased from Dharmacon. HeLa or mouse 3T3 cells were transfected using a mixer of Lipofectamine RNAiMax (Invitrogen) and siRNA oligos (5-10 nM) and analyzed at 48-72 hrs after transfection. The siRNA sequences used in this study are as follow, siTopIIA-04, CGAAAGGAAUGGUUAACUA; siTopIIA #18, AGUGACAGGUGGUCGAAAU; siTopIIB-06, GAACUUUCCUUCUACAGUA (Dharmacon, D-004240-06-0002); siTopIIB #19, GAUGAUAGUUCCUCCGAUU (Dharmacon, D-004240-19-0002); siTopI #1, GAAAGGAAAUGACUAAUGA (Dharmacon, D-005278-01-0002); siTopI #2, GAAGAAGGCUGUUCAGAGA (Dharmacon, D-005278-02-0002); siSgo1 GAGGGGACCCUUUUACAGATT; siMre11#1, GAUGAGAACUCUUGGUUUA (Dharmacon D-009271-01-0002); siMre11#4, GAGUAUAGAUUUAGCAGAA (Dharmacon D-009271-01-0002); simTopI#1, GCACUGUACUUCAUUGAUA (Dharmacon D-047567-01-0002); simTopI #4, UAGCAAAGACGCAAAGGUU (Dharmacon D-047567-04-0002).

### Antibodies and Immunoblotting

Antibodies used in this study: anti-centromere antibody (ACA or CREST-ImmunoVision, HCT-0100), anti-Smc1 (Bethyl, A300-055A), anti-Rpb1 (Abcam, ab5408), anti-Actin (Invitrogen, MA5-11869), anti-Rpb1-pSer2 (Biolegend, H5), anti-Rpb1-pSer2 (Active motif, 61083), anti-γH2AX (Cell signaling, 2577), anti-pATM^S1981^ (Rockland, 200-301-400), anti-Top 1 (Bethyl, A302-589A), anti-TopIIa (Bethyl, A300-054A), anti-TopIIb (Bethl, A300-949A), anti-Aurora B (BD Biosciences, 611083), anti-Rpb (abcam, 4H8, ab5408), anti-α-Tubulin (Abcam, ab4074), anti-ATM (Novus,

NB100-309), anti-TopI-cc (MilliporeSigma, MABE1084), anti-DNA-PK-pS2056 (Thermo, PA5-78130), anti-CENP-A (EMD Millipore; 07–574), anti-CENP-B (Abcam; ab25734), anti-CENP-C (MBL, PD030), anti-RNAP II (8W1G16, Abcam), anti-Smc1 (Bethyl; A300-055A), anti-γH2Av (DHSB, UNC93-5.2.1). Anti-Sgo1, anti-Knl1 and anti-Bub1 were made in-house as described previously ^54–56^. Anti-Sororin antibodies described previously were a gift from Dr. Susannah Rankin ^57^.

The secondary antibodies were purchased from *Li-COR*: IRDye^®^ 680RD Goat anti-Mouse IgG Secondary Antibody (926-68070) and Goat anti-Rabbit IgG Secondary Antibody (926-32211). For immunoblotting, primary and secondary antibodies were used at 1 μg/ml concentration.

### CRISPR/Cas9

DNA oligos that target α-satellite DNAs or genes (CSB) were cloned to px330 (U6). Sequence-validated plasmids were transfected into cells using Effectene transfection reagent (Qiagen) for 12 or 24 hrs. The gRNA sequence for α-satellite DNAs was previously described^58^ and the one for CSB was the following: GCAGAAGTCGGAGTCGCTGT.

### RNA extraction, Reverse transcription, and real-time PCR analysis

Cells or tissues were then collected and then dissolved in TRIzol solution (Invitrogen, 15596026) for RNA extraction. Extracted total RNAs were finally dissolved in nuclease-free water and treated with TURBO DNase (Invitrogen, AM2238) in the presence of RNase inhibitor (NEB, M3014) at 37°C for 45 min. After being extracted with Phenol/ Chloroform/Isoamyl alcohol (25:24:1, v/v) (Invitrogen, 15593-031) and precipitated with ice-cold ethanol solution containing glycogen and sodium acetate, total RNAs were finally dissolved in nuclease-free water.

Purified RNAs from cells and tissues were mixed with iScript Reverse Transcription Supermix (Bio-Rad, 1708841) and reverse transcription was performed according to the manufacturer’s protocols. After being mixed with the SsoAdvanced™ Universal SYBR® Green Supermix (Bio-Rad, 1725274), the synthesized cDNA was subjected to real-time PCR analysis using QuantStudio 6 Flex Real-Time PCR System (Applied Biosystems).

The primers for human cells were used in this study: GAPDH-F TGATGACATCAAGAAGGTGGTGAAG, GAPDH-R TCCTTGGAGGCCATGTGGGCCAT; Rpl30-FCAAGGCAAAGCGAAATTGGT, Rpl30-R GCCCGTTCAGTCTCTTCGATT; SAT-1-F AAGGTCAATGGCAGAAAAGAA, SAT-1-R CAACGAAGGCCACAAGATGTC; SAT-4-F CATTCTCAGAAACTTCTTTGTGATGTG, SAT-4-R CTTCTGTCTAGTTTTTATGTGAATATA; SAT13/21-F TAGACAGAAGCATTCTCAGAAACT, SAT-13/21-R TCCCGCTTCCAACGAAATCCTCCAAAC. Primers for different α-satellite HORs used in extended data fig.4a were described previously^59^. Mre11-F GTCCGTGAGGCTATGACCAG, Mre11-R CAGACCAGTGTCTGCTCTTCC. Ribosomal DNA 18S-F, CTCAACACGGGAAACCTCAC, Ribosomal DNA 18S-R, CGCTCCACCAACTAAGAACG; Ribosomal DNA 28S-F, GACCCGAAAGATGGTGAACT, Ribosomal DNA 28S-R, CCGGGCTTCTTACCCATTTA.

The primers for mouse cells were used in this study. mGAPDH, AGGTCGGTGTGAACGGATTTG and TGTAGACCATGTAGTTGAGGTCA; mRpl30, TTGGTACAGAATGGATTCGTCAC and GGGTCCCCACCATACTTTTCA; minor satellite, CATGGAAAATGATAAAAACC and CATCTAATATGTTCTACAGTGTGG; major satellite, GACGACTTGAAAAATGACGAAATC and CATATTCCAGGTCCTTCAGTGTGC. mTopI, GACCATCTCCACAACGATTCC and ATGCCGGTGTTCTCGATCTTT.

The primers for *Drosophila* used in this study were described previously ^40^. TopI-F, TGTAACCATCAGCGTTCCGT; TopI-R, TTCAGCTGATCCCTTAGGCG; CEN359-R, TATTCTTACATCTATGTGACC; CEN359-L, GTTTTGAGCAGCTAATTACC; Rp49, ATGACCATCCGCCCAGCATAC and CTGCATGAGCAGGACCTCCAG; PGd, AGGACTCGTGGCGCGAGGTG and GGAATGTGTGAACGGGAAAGTGGAG; dGAPDH-F, TAAATTCGACTCGACTCACGGT; dGAPDH-R, CTCCACCACATACTCGGCTC.

### EU chasing and purification of EU-RNAs

EU-RNAs were prepared according to the protocol from Click-iT™ Nascent RNA Capture Kit (C10365, ThermoFisher). Cells with a confluency of 60–80% in 10 cm petri dishes were incubated with EU at a final concentration of 0.5 mM for 1 hr. Collected EU-treated cells were then dissolved in TRIzol solution (Invitrogen, 15596026). Total RNAs were extracted according to the section of “RNA extraction”. Extracted total RNAs were finally dissolved in nuclease-free water and treated with TURBO DNase (Invitrogen, AM2238) in the presence of RNase inhibitor (NEB, M3014) at 37°C for 45 min. After being extracted with Phenol/ Chloroform/Isoamyl alcohol (25:24:1, v/v) (Invitrogen, 15593-031) and precipitated with ice-cold ethanol solution containing glycogen and sodium acetate, total RNAs were finally dissolved in nuclease-free water. These RNAs were then further incubated with streptavidin dynabeads pretreated with Salmon sperm DNA (Invitrogen, 15632-011) in binding buffer for 45 min. With the help of DynaMag™-2 Magnet (Invitrogen, 12321D), dynabeads were washed with wash buffer I and II. The bead-captured EU-RNAs were converted to cDNA using iScript DNA synthesis kit (Bio-Rad) and further subjected to real-time PCR analysis.

### Flow cytometry

Cultured cells that were harvested cells by trypsinization were washed with PBS (PH7.4), and fixed with ice-cold 70% ethanol overnight at −20°C. After ethanol was washed out with PBS, cells were further permeabilized with PBS (PH 7.4) containing 0.25% Triton X-100 for 5 min. Finally, cells were stained with propidium iodide (Sigma-Aldrich) at a final concentration of 20 µg/ml. RNase A (QIAGEN) was added at a final concentration of 200 µg/ml. The samples were analyzed with BD LSR Fortessa flow cytometer. The data were analyzed by software Modfit.

### Comet assay

Cells were then mixed with comet agarose at a ratio of 1:10 (v/v) after resuspended at a density of 1X10^5^ cells/ml with PBS. A drop of the mixer was placed onto an agarose-coated slide, flattened, and covered by a cover glass. After horizontally kept at 4°C for 15 mins, cover glass was gently removed. The slide was sequentially immersed at 4°C in the dark with cold lysis buffer (2.5M NaCl, 100mM EDTA, 1% DMSO, 1X Lysis solution) for 45 mins, cold alkaline buffer (0.3M NaOH, 1mM EDTA) for 30 mins, and cold TBE buffer for 5 mins. Then, the sidle was transferred into electrophoresis chamber filled with cold TBE electrophoresis buffer and run at 1 V/cm for 10-15 mins. After immersed with cold water for 2 mins twice and with cold 70% ethanol for 5 min twice, the slide was stained with Vista Green DNA Dye at room temperature for 15 mins. The slide was sealed with cover glass and then subjected to microscopic analysis.

### Manipulation of Drosophila larvae and solid tumors

Drosophila lines were maintained and crossed at room temperature (25°C) on the BDSC cornmeal food (https://bdsc.indiana.edu/information/recipes/bloomfood.html). The following fly lines were used in this study: Act-Gal4/CyO; tub-Gal80ts/TM6B, UAS-GFP, UAS-top1-RNAi (Bloomington stock center: 35424#3, dsRNA-GL00347: CACCAAGGAAGTGTTCAATAA; 55314#5, dsRNA-HMC04001: CTGCACCAAGGAAGTGTTCAA), and UAS-NICD ^42,43^. The inducible *Gal4/Gal80^ts^* system with *Act-Gal4* was used to ubiquitously overexpress genes or RNAi in cells. For these experiments with Gal80^ts^, eggs expressing *UAS-RFP TubGal80^ts^/+; Act-Gal4* with or without *UAS-top1-RNAi* were raised at 18°C (Gal4 is ‘off’) for 2 days and then shifted to 29°C to degrade Gal80^ts^ (Gal4 is ‘on’) for indicated days so that *Act-Gal4* can drive expression of *UAS-top1-RNAi*. After that, the wing imaginal discs were used for RNA extraction. Induction of *Drosophila* larval salivary gland imaginal ring tumor was previously described in ^42,43^. Flies with *Act-Gal4/UAS-NICD; tub-Gal80^ts^/+* were raised at 18°C to inhibit Gal4 function until late-second instar and then were transferred to 29°C to induce NICD expression. Flies were allowed to lay eggs at 18°C for one day and reared at 18°C for 7 days. After 7 days, late-second instar larval fly were transferred to 29°C for GFP and NICD induction. The control fly became pupal and adult after 3 days at 29°C. NICD induced fly showed developmental delay during the larval stage and most of them died after 7 days at 29℃.

As for *Drosophila lethal (2) giant larvae (lgl)* tumor, Fly lines were used as follows: w; lgl4 FRT40A/CyO, yw; Act-Gal4,tub-Gal80ts (Actts-Gal4)/TM6B, UAS-Top1RNAi. 1-day eggs were collected at 18°C, and eggs were cultured at 18°C for 7 days, then shifted to 29°C for additional days as indicated. Wing discs from giant larvae w; lgl4 FRT40A/lgl4 FRT40A, and yw; lgl4 FRT40A/lgl4 FRT40A; Actts-Gal4/UAS-Top1RNAi were dissected and used for RNA extraction.

RNA extraction, cDNA synthesis and quantitative real-time PCR were performed as follows. Total RNAs were extracted from eighty tumor ImRs or control ImRs from larvae by using Zymo RNA preparation kit (Cat. No.: R2070) according to manufacturer’s instructions. DNA was removed by treating with RNase Free DNase Set. cDNA synthesis was performed using the Roche First Strand cDNA synthesis Kit (53759220) according to manufacturer’s instructions. Quantitative real time PCR was performed using SYBR Green Master Mix (BIO-RAD, #1725121) with gene-specific primer sets, on a C1000 Touch Thermal Cycler (CFX96, BIO-RAD). Comparative qPCRs were performed in triplicate using the above primers.

### Chromatin immunoprecipitation (ChIP)

HeLa cells were crosslinked with buffer (50 mM Hepes PH 8.0, 1% Formaldehyde, 100 mM NaCl, 1 mM EDTA, 0.5 mM EGTA) at room temperature for 10 min and further treated with 125mM glycine for another 5 min. Cells were then resuspended in IP buffer (10 mM Tris, 300 mM NaCl, 1 mM EDTA, 1 mM EGTA, 1% Triton X-100, 1% Sodium deoxycholate) and sonicated using a Fisher Scientific sonicator. After cell debris was removed by centrifugation, the supernatant was incubated with protein-A beads (Santa Cruz, SC-2001) at 4°C for 2 hrs. Pre-cleared cell lysate was incubated with 5 µg antibodies overnight and further with protein-A beads for another 2 hrs at 4°C. Pelleted beads were sequentially washed by low salt buffer (20 mM Tris 8.0, 150 mM NaCl, 0.1% SDS, 1% Triton X-100, 2 mM EDTA), high salt buffer (20 mM Tris 8.0, 500 mM NaCl, 0.1% SDS, 1% Triton X-100, 2 mM EDTA), LiCl buffer(10 mM Tris 8.0, 0.25 M LiCl, 1% IGEPAL CA630, 1% sodium deoxycholate, 1 mM EDTA), and TE buffer (10 mM Tris PH 8.0, 1 mM EDTA PH 8.0). Washed beads were treated with elution buffer (10 mM Tris 8.0, 1 mM EDTA, 1% SDS) at 65°C for 10 min and the resulting supernatant was further incubated at 65°C overnight to reverse the crosslinking. Then the solution was sequentially treated with RNase A (Qiagen 1007885) at 37°C for 1 hr and Proteinase K (Thermo Fisher Scientific EO0491) at 50°C for 2 hrs. Finally, DNA in the solution was extracted with Phenol/ Chloroform/Isoamyl alcohol (25:24:1, v/v) (Invitrogen, 15593-031) and purified by Qiagen gel purification kit for real-time PCR analyses.

### Coimmunoprecipitation

Cells with different treatments were dissolved in lysis buffer (25 mM Tris-HCl at pH 7.5, 50 mM NaCl, 5 mM MgCl_2_, 0.1% NP-40, 1 mM DTT, 0.5 μM okadaic acid, 5 mM NaF, 0.3 mM Na_3_VO_4_ and 100 units/ml TurboNuclease (Accelagen)). After an incubation of 1 hr on ice followed by another 10-min incubation at 37 °C, the lysate was cleared by centrifugation for 15 min at 4 °C at 20,817g. The resulting supernatant was incubated with the antibody overnight at 4 °C. In the next day, protein-A beads were added and further incubated for another 1 hr before washed four times with wash buffer (25 mM Tris-HCl at pH 7.5, 50 mM NaCl, 5 mM MgCl_2_, 0.1% NP-40, 1 mM DTT, 0.5 μM okadaic acid, 5 mM NaF, and 0.3 mM Na_3_VO_4_). The proteins bound to the beads were finally dissolved in SDS sample buffer, separated by SDS-PAGE and blotted with the appropriate antibodies.

### Immunofluorescence and chromosome spread

Chromosome spread and immunostaining were performed as described before ^55^. Nocodazole-arrested mitotic cells were swollen in a hypotonic solution containing 50 mM KCl for 15 min at 37°C and then spun onto slides with a Shandon Cytospin centrifuge. Cells were then sequentially treated with ice-cold PBS containing 0.2% Triton X-100 for 2 min and with 4% ice-cold paraformaldehyde for 4 min. After being washed with PBS containing 0.1% Triton X-100, cells were incubated with primary antibodies overnight at 4°C. Cells were then washed with PBS containing 0.1% Triton X-100 and incubated at room temperature for 1hr with the appropriate secondary antibodies conjugated to fluorophores (Invitrogen, A11008, A21090 and A31571). After being washed again with PBS containing 0.1% Triton X-100, cells were stained with 1 μg/ ml DAPI and mounted with Vectashield. The images were taken by a Nikon inverted confocal microscope (Eclipse Ti2, NIS-Elements software) with a ×60 objective. Image processing was carried out with ImageJ and Adobe Photoshop. Quantification was carried out with ImageJ. Statistical analysis was performed with Graphpad Prism.

### Chromosome Stretching

Cell, after treated with salt detergent buffer (25 mM Tris pH 7.5, 500m M NaCl, 1% Triton in water), were spun onto slides at 1000 rpm for 4 min with a Shandon Cytospin centrifuge. Slides were further placed into salt detergent buffer for 10 mins and then gently and slowly taken out. These slides were then further subjected to regular immunostaining.

### RNA-FISH

Cells were spun onto slides pre-treated with RNase away. After being treated ice-cold PBS containing 0.2% Trition X-100 for 2 mins and 4% ice-cold paraformaldehyde for 4 mins or, cells on the slide were washed with 2X SSC buffer twice and further incubated with 70% ethanol at 4°C overnight. Alternatively, Cells on slides were fixed with 4% room-temperature paraformaldehyde (in PBS) for 10 mins. After being washed twice with PBS, cells were permeabilized in 70% ethanol for 20 mins. Cells with either of above treatments were finally hybridized with probes coupled with FAM (Qiagen, probe-1, TTCTGAGAATGCTTCTGTCTA; probe-2, ACGTCCGCTTGCAGATACTACA) diluted in hybridization buffer at 37°C overnight. After being washed once with washing buffer at 37°C for 30 mins and three times with 2X SSC buffer (in the last time, DAPI was added), Cells on slides were ready for microscopic analysis. For FISH and ACA co-staining, cells were firstly treated with the RNA-FISH procedures described above and then incubated with ACA antibody and secondary antibody. After being stained with DAPI, cells are ready for imaging.

### RNA-sequencing (RNA-seq) analysis

Paired-end strand-specific ribodepleted total RNA-seq reads were first validated by the FastQC algorithm. Raw sequence reads were then aligned to a complete Telomere-to-Telomere (T2T) reconstructed human reference genome (T2T-CHM13 v1.0)^21^. The alignments were performed using Spliced Transcripts Alignment to a Reference (STAR) aligner version 2.5.3a^60^ and were subjected to visual inspection using the Integrative Genomics Viewer (IGV) genome browser^61^. Transcript data from STAR were subsequently analyzed using RSEM version 1.3.0^62^ for quantification of human centromere transcripts. Read coverage data were generated using the bamCoverage tool and visualized using the IGV genome browser as previously described^63^.

To determine how CPT and etoposide treatments globally affect gene expression in RPE-1 cells, the False Discovery Rate (FDR) was calculated between three pairs of samples, DMSO/CPT, DMSO/etoposide, and DMSO/etoposide plus CPT, using the EBSeq algorithm^64^ to identify the differentially expressed genes. Based on values of FDR less than 0.05, CPT, etoposide, and etoposide plus CPT treatments significantly changed the expression of 1079, 539 and 1304 genes, respectively. These genes only account for 1.7%, 0.87% and 2.1% of total human genes (∼63,000).

To determine how TopI inhibition affects the expression of lowly transcribed genes (extended data fig.6c). we firstly defined a group of lowly transcribed genes, in which, each member had a TPM (Transcripts Per Million) of no more than 2, as opposed to an average TPM of 16 per gene globally. Out of ∼62,000 genes, about 10% (∼7,800 genes) fell into this group. CPT treatment for 12 hrs affected the expression of only ∼10% of genes (∼780 genes) in this group. Volcano analysis on gene expression in this group treated with DMSO or CPT was shown in extended data Figure S9C.

In figs.1e,3f, TPM was firstly obtained for each annotated HOR using a complete Telomere-to-Telomere reconstructed human reference genome^21^. Average of fold changes for HORs with detected transcripts was then calculated in CPT treated cells.

### Quantification and Statistical analysis

Numeric values for the intensities of experimental subjects under investigation were obtained with Image J. As for quantification in Figures 2C and S1E, 5-6 centromeres were randomly selected from each cell, a mask was generated to mark centromeres based on ACA fluorescence signals in the projected image. After background subtraction, the intensities of RNAP II (Rpb1), TopI-cc, and ACA signals within the mask were obtained in number. Relative intensity was calculated from the intensity of RNAP II (Rpb1) signals normalized to the one of ACA signals and plotted with the GraphPad Prizm software. All the experiments were repeated at least three times.

For quantification in Figures 3H, 3I, 4H, 4I, S1F, S4B, S6A, a mask was generated to mark nuclei based on DAPI signals in the projected image. After background subtraction, the intensity of γH2AX or DAPI fluorescence signals within the mask were obtained in number. Relative intensity was calculated from the intensity of γH2AX signals normalized to the one of DAPI signals and plotted with the GraphPad Prizm software. All the experiments were repeated at least three times.

For quantification in Figure 2B and S6B, a mask was generated to mark the stretched centromeric chromatin based on DAPI and ACA signals in the projected image. After background subtraction, the intensity of γH2AX, RNAP II-pSer2 or DAPI fluorescence signals within the mask were obtained in number. Relative intensity was calculated from the intensity of γH2AX, or RNAP II-pSer2 signals normalized to the one of DAPI signals and plotted with the GraphPad Prizm software. All the experiments were repeated at least three times.

Quantification was usually performed based on the results from at least three independent experiments. All the samples analyzed were included in quantification. Sample size in figures was recorded in source files. No specific statistical methods were used to estimate sample size. No methods were used to determine whether the data met assumptions of the statistical approach.

Differences were assessed using ANOVA followed by pairwise comparisons using Tukey’s test (Figures 1G, 3H, 3I, 4D, 4E, 5C, 4E, 5G, 5H, S4B, S4C, S8C, S8E, S8F) or two-sided Student’s T-test (Figures 1A,1B,1C,1D, 1F, 2A, 2B, 2C, 3A, 3B, 3D, 3E, 3G, 4A, 4B, 4C, 4H, 4I, 5A, 5B, 5D, 5F, S1A, S1E, S1F, S3A,S3B, S3C, S3E, S4D, S4E, S6A, S6B, S6C, S6D, S6E, S6F, S6G, S7A, S8A, S8D, S8F, S8G).

## Supplemental Figures

**Figure S1.**
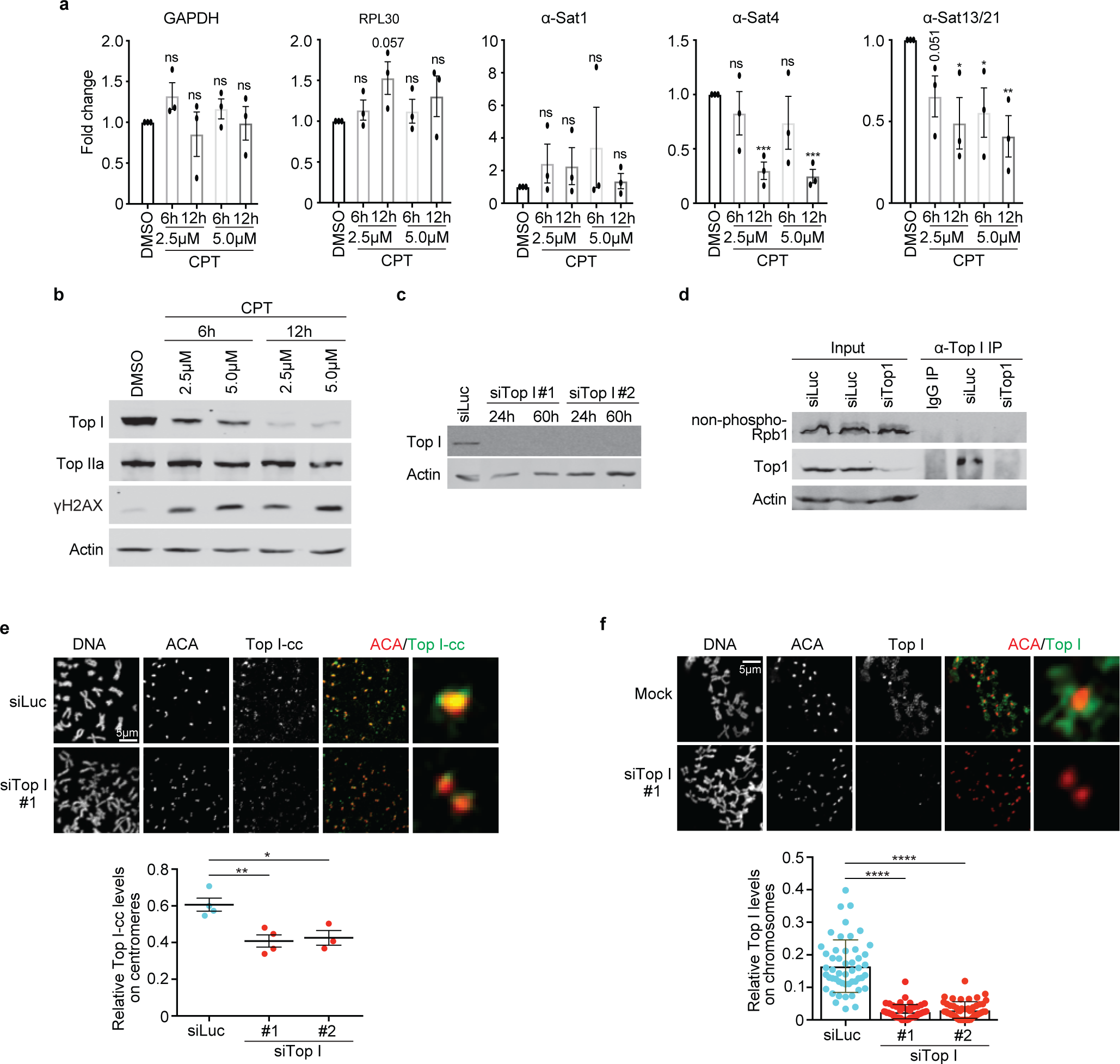
TopI regulates α-satellite transcription A. TopI inhibition decreases centromeric transcription. HeLa cells were treated with CPT (2.5 μM) at distinct doses for 6 or 12 hrs and RNAs were extracted for real-time PCR analysis. **B.** Lysates of HeLa cells in (**A**) were subjected to Western-blotting analysis. **C. Knockdown efficiency of TopI siRNA.** Lysates of HeLa cells transfected with luciferase or TopI siRNAs (#1 or #2) at different timepoints were subjected to Western-blotting analysis. **D. TopI does not physically interact with non-phosphorylated RNAP II**. Lysates of HeLa cells transfected with luciferase siRNA or TopI siRNA#1 were subjected to coimmunoprecipitation with antibody against non-phosphorylated RNAP II (8WG16). Results for coimmunoprecipitation with antibodies against total RNAP II (4H8) and phosphorylated RNAP II (Ser2) (H5) were recorded in Figure 2D. **e** and **f. Validation of and TopI and TopI-cc fluorescence signals.** HeLa cells were transfected with luciferase siRNA or TopI siRNA#1. After a brief nocodazole (1.5 hr, 5 μM) treatment, mitotic cells were collected for immunostaining. Quantifications of TopI-cc levels (TopI-cc/ACA, **E**) and TopI levels (TopI/DNA, **F**) were shown in the bottom panels.

**Figure S2.**
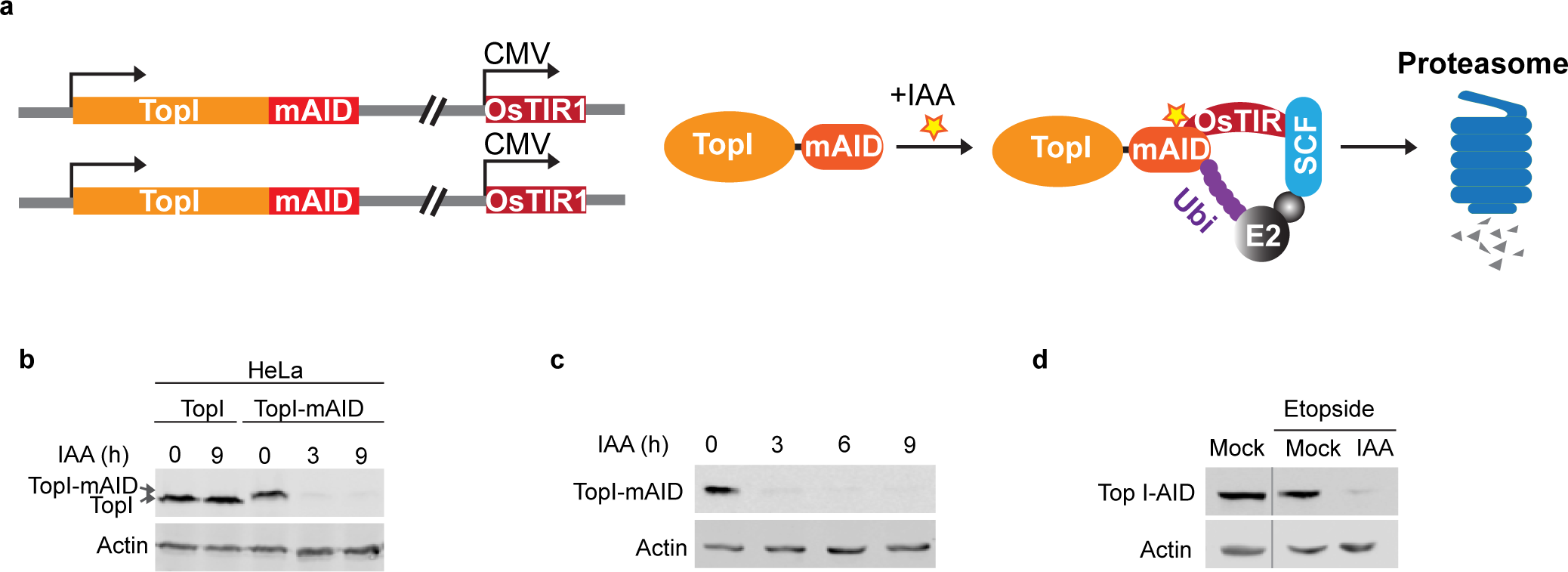
Construction of TopI-mAID cell lines. **A.** Schematic for construction of TopI-mAID**. B.** Parent HeLa cells and Top-mAID HeLa cells were treated with IAA (1 μM) at the indicated times. Cell lysates were analyzed by Western blots. **C.** Top-mAID HeLa cells were treated IAA at the indicated times. Cells were collected for real-time PCR analysis (recorded Fig. 1d) and Western blotting analysis. **D.** Top-mAID HeLa cells were treated IAA (1 μM) and etoposide for 9 hrs. Cells were collected real-time PCR analysis (recorded in Fig. 3d) and Western blotting analysis.

**Figure S3.**
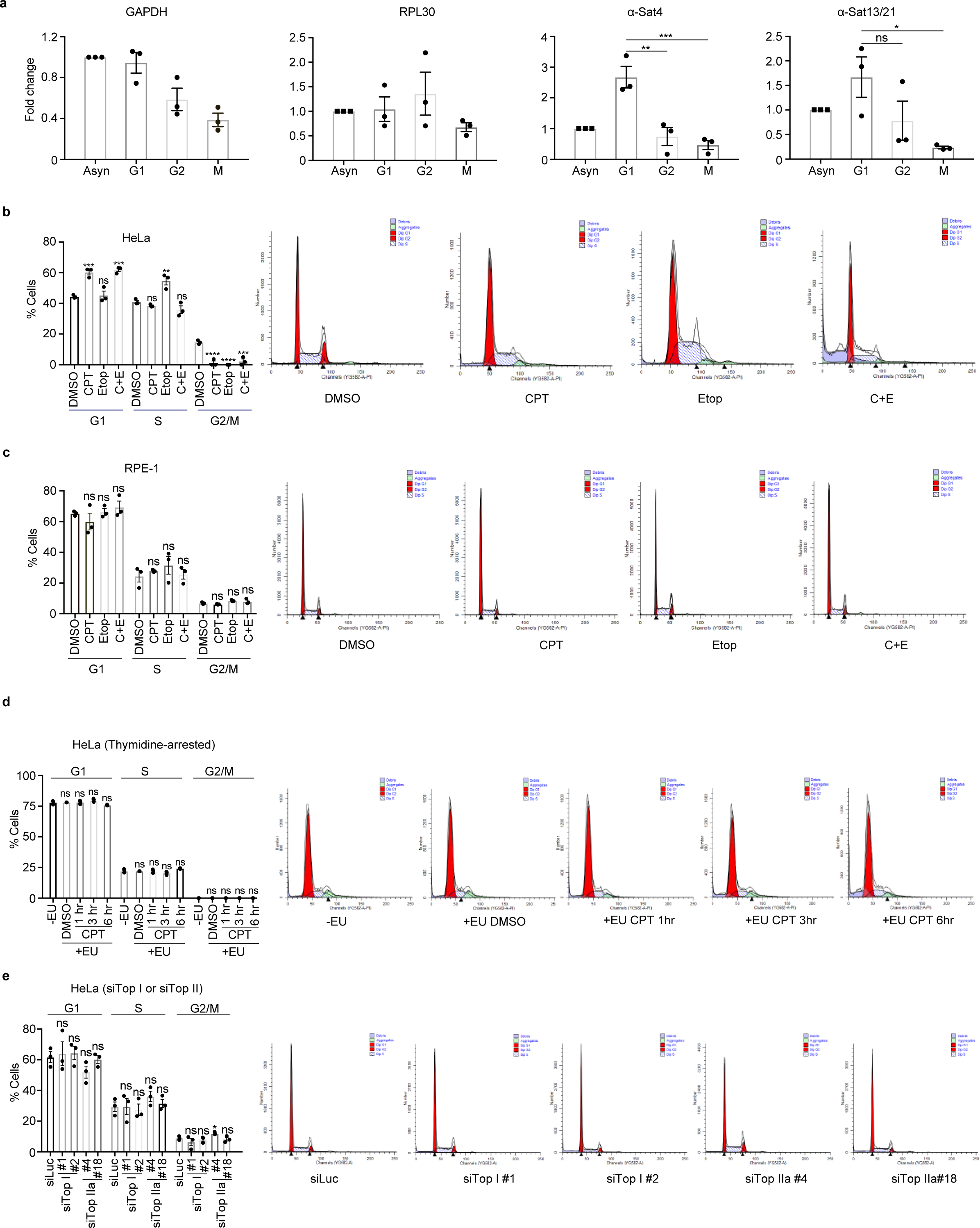
Flow-cytometric analyses of HeLa cells. >**A. α-satellite transcription is cell cycle-regulated.** RNAs extracted for HeLa cells arrested with thymidine (18 hrs, G1), RO3306 (18 hrs, G2) or nocodazole (16 hrs, M), were subjected to real-time PCR analysis. **B** and **C**. **Flow-cytometric analyses of HeLa cells with chemical treatments.** HeLa (**B**) or RPE-1 (**C**) cells treated with DMSO, CPT (2.5 μM), Etoposide (Etop, 30 μM), or CPT plus Etoposide (C+E) for 12 hrs, were analyzed with FACS. Quantifications of cell cycle profiles were shown in the left panel. Representatives of FACS results were shown in the right panels. **D. Flow-cytometric analyses of Thymidine-arrested HeLa cells with CPT treatment at different timepoints.** HeLa cells were incubated with thymidine for 18 hrs, during which, cells were also treated with EU and/or CPT (2.5 μM) at different timepoints. Cells were then collected for FACS analysis. **E. Flow-cytometric analyses of HeLa cells transfected with TopI or TopII siRNAs.** HeLa cells transfected with luciferase, TopI or TopII siRNAs for 48 hrs, were collected for FACS analysis.

**Figure S4.**
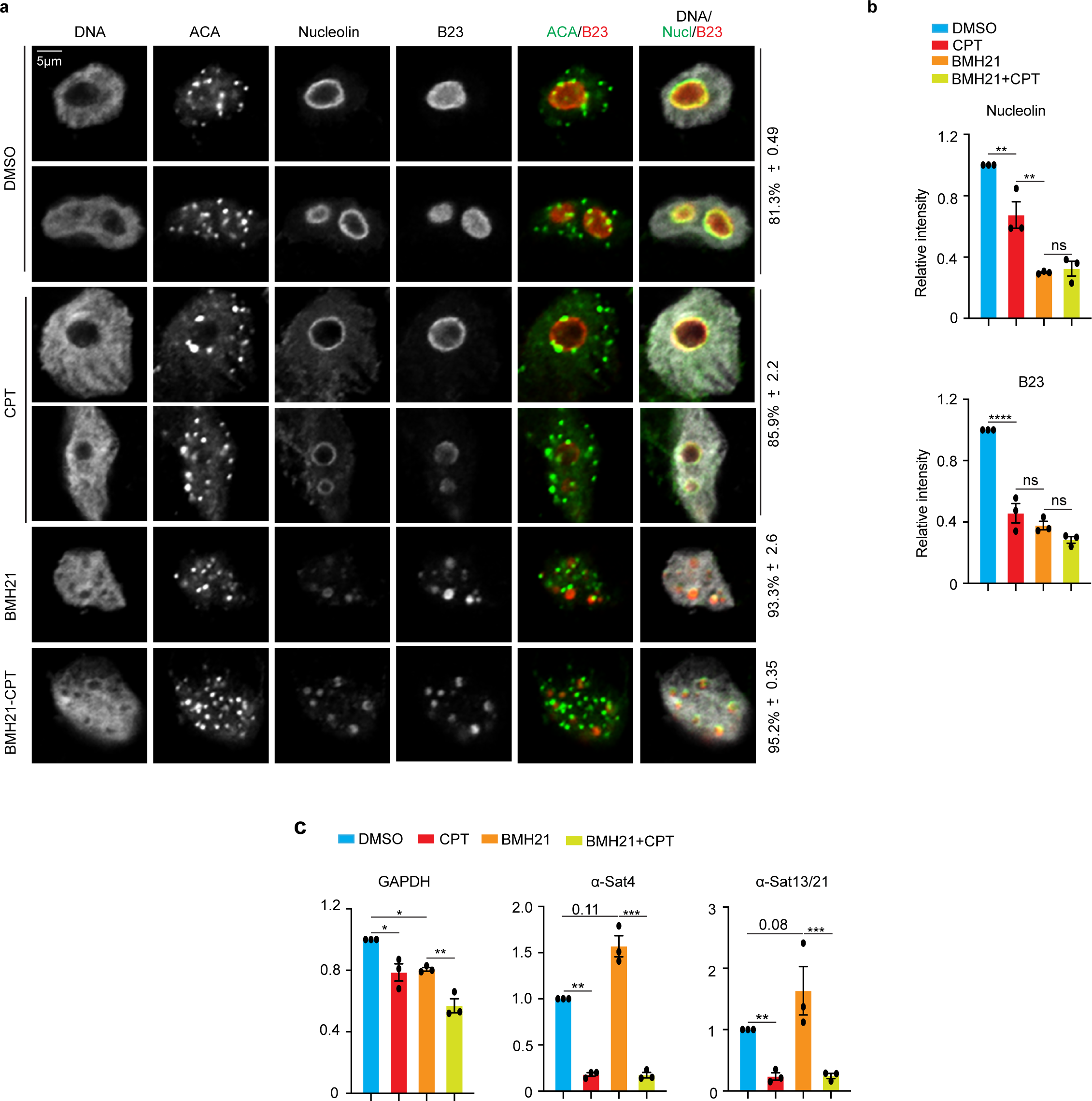
The nucleolus may not play an important role in TopI-mediated α-satellite transcription. **A. RNAP I inhibitor BMH-21 treatment disrupts the nucleolar structure and decreases the levels of nucleolin and B23.** HeLa cells treated with DMSO, CPT (10 μM), BMH-21 (RNAPI inhibitor), or BMH-21 plus CPT for 12 hr were stained with DAPI and the indicated antibodies. **B.** Relative intensities of nucleolar components nucleolin (nucleolin/DNA) and B23 (B23/DNA) in (**A**) were shown here. **C**. **TopI-regulated α-satellite transcription is independent of intact nucleolar structures.** HeLa cells treated as described in (**A**) were collected for real-time PCR analysis.

**Figure S5.**
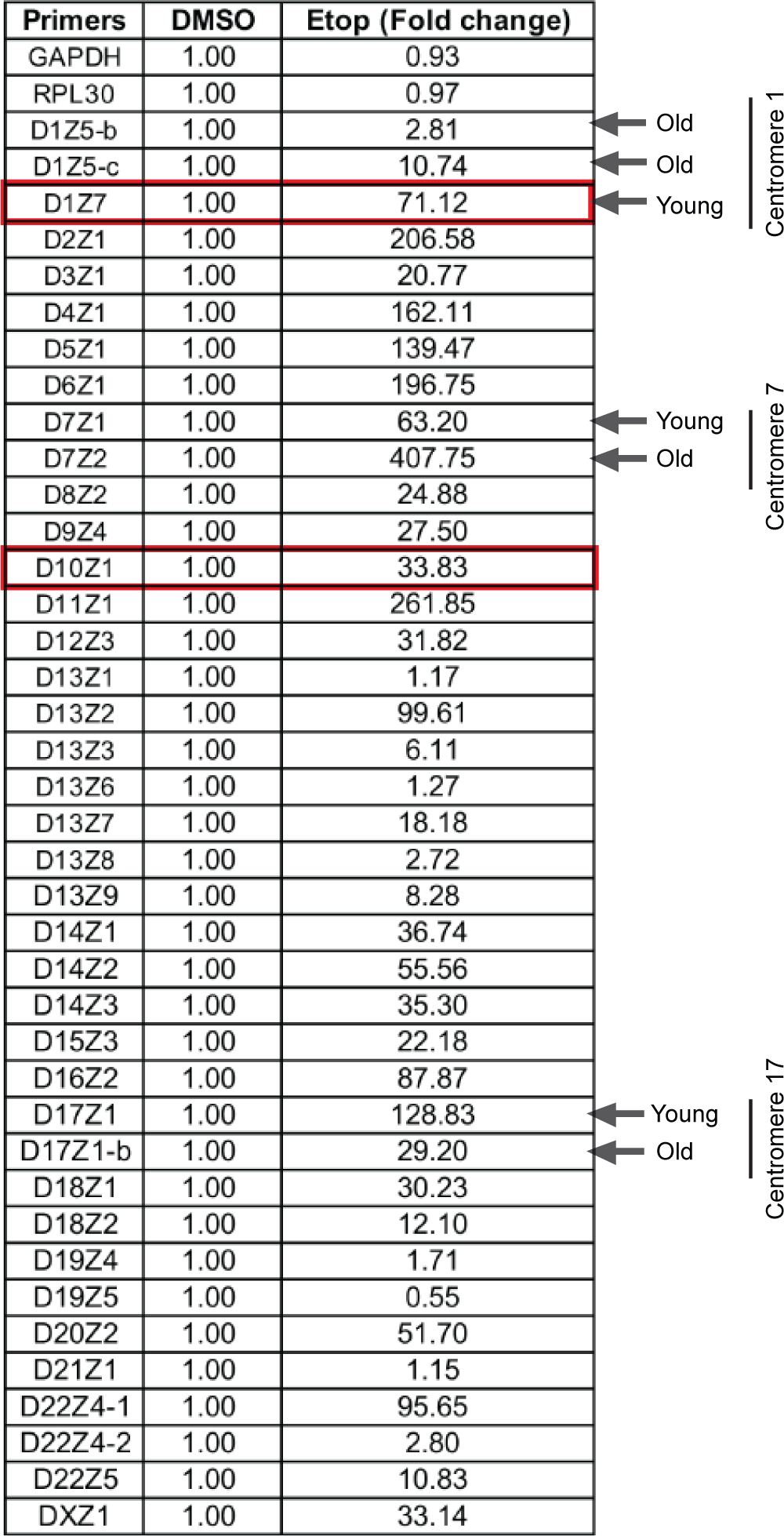
Real-time PCR CT values for different kinds of α-satellite arrays and etoposide globally induces α-satellite transcription. **Real-time PCR analyses of centromere RNAs using primers against distinct types of α-satellite HORs.** RNAs extracted from etoposide-treated HeLa cells (30 μM, 12 hrs) were subjected to real-time PCR analysis using primers against different types of. α-satellite HORs. Two boxed regions in red are α-satellite HORs for centromeres 1 and 20. Young and old denote youngest α-satellite HOR and oldest α-satellite HOR, as described in^20^.

**Figure S6.**
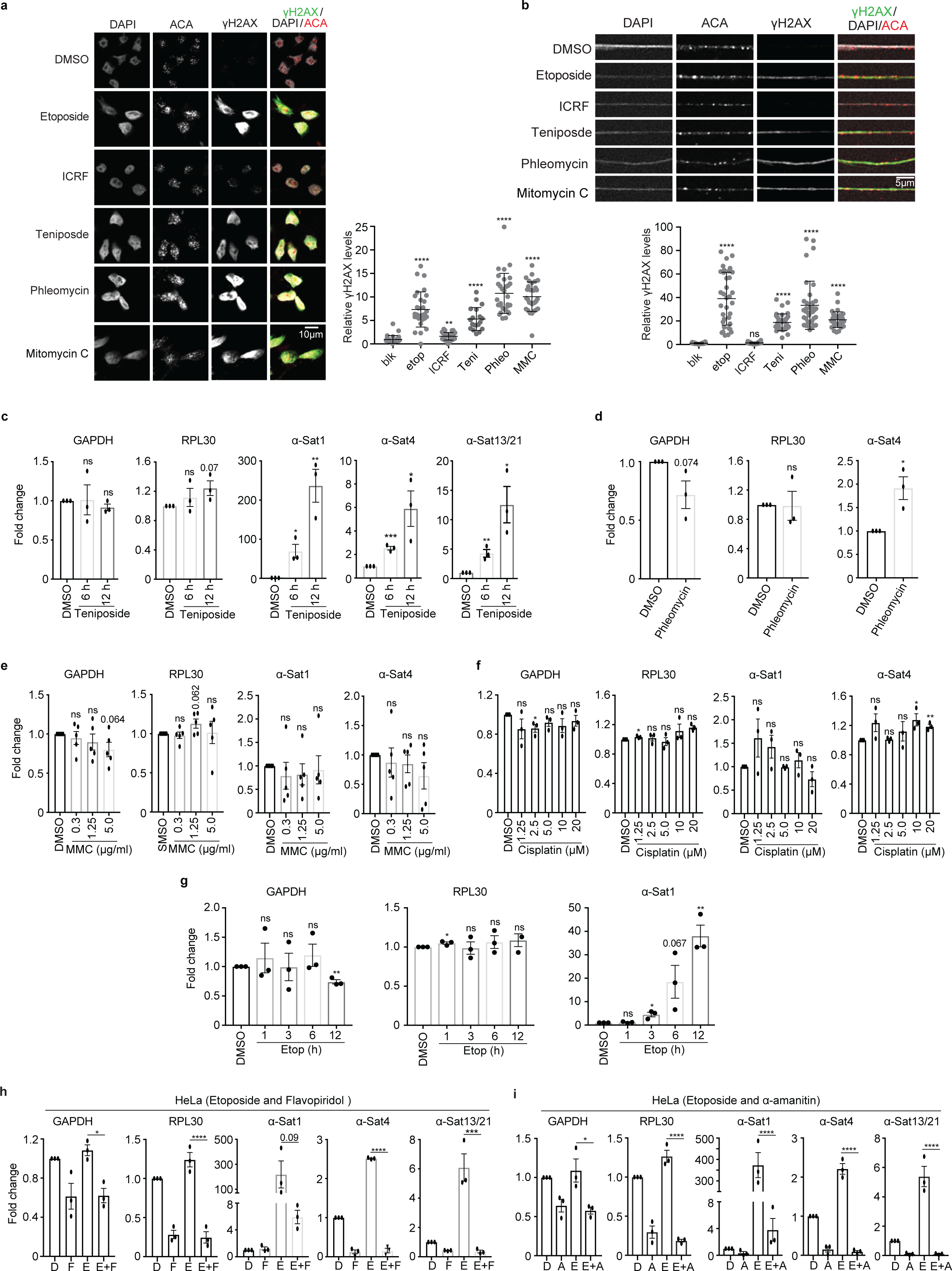
DSBs dramatically increases α-satellite transcription. **A. DNA-damage analyses of HeLa cells with chemical treatments.** HeLa cells were treated with DMSO, etoposide (Etop, 30 μM), ICRF (20 μM), teniposide (Teni, 2.5 μM), phleomycin (phleo, 80 μg/ml), or mitomycin C (MMC, 5 μg/ml) for 12 hrs, and then subjected to immunostaining. Quantitation of γH2AX levels (γH2AX/DNA) was shown in the right panel. **B. DNA-damage analyses of centromeres in HeLa cells with chemical treatments**. HeLa cells treated with DMSO, etoposide (Etop) (12 hrs, 30 μM), ICRF (20 μM), teniposide (Teni) (12 hrs, 2.5 μM), phleomycin (Phleo) (24 hrs, 80 μg/ml), or mitomycin C (MMC) (12 hrs, 5 μg/ml), were subjected to chromatin stretch followed by immunostaining. Quantitation or γH2AX levels (γH2AX/DNA) on centromeric regions was shown in the bottom panel. **C**, **D**, **E**, and **F, Effects of various chemicals on α-satellite transcription.** HeLa cells were treated with DMSO, teniposide (**c**) (12 hrs, 2.5 μM), phleomycin (**D**) (24 hrs, 80 μg/ml), mitomycin c (**E**) (12 hrs, indicated concentrations), or cisplatin (**F**) (12 hrs, indicated concentrations), and RNAs were extracted for real-time PCR analysis. Results for α-Sat13/21 primers were shown in fig.2d. **G. Etoposide increases α-satellite transcription in a time-dependent manner.** HeLa cells were treated with DMSO or etoposide (2.5 μM) at the different timepoints and RNAs were extracted for real-time PCR analysis. Results from α-Sat13/21 primers were recorded in fig.2g. **H** and **I. DSB-induced α-satellite transcription depends on RNAP II.** HeLa cells treated with RNAP II inhibitor α-amanitin (**i,** 50 μg/ml), flavopiridol (**H,** 1 μM), etoposide (**H** and **I**, 30 μM), flavopiridol plus etoposide (**H**), or α-amanitin plus etoposide (**I**) for 12 hrs and RNAs were extracted for real-time PCR analysis.

**Figure S7.**
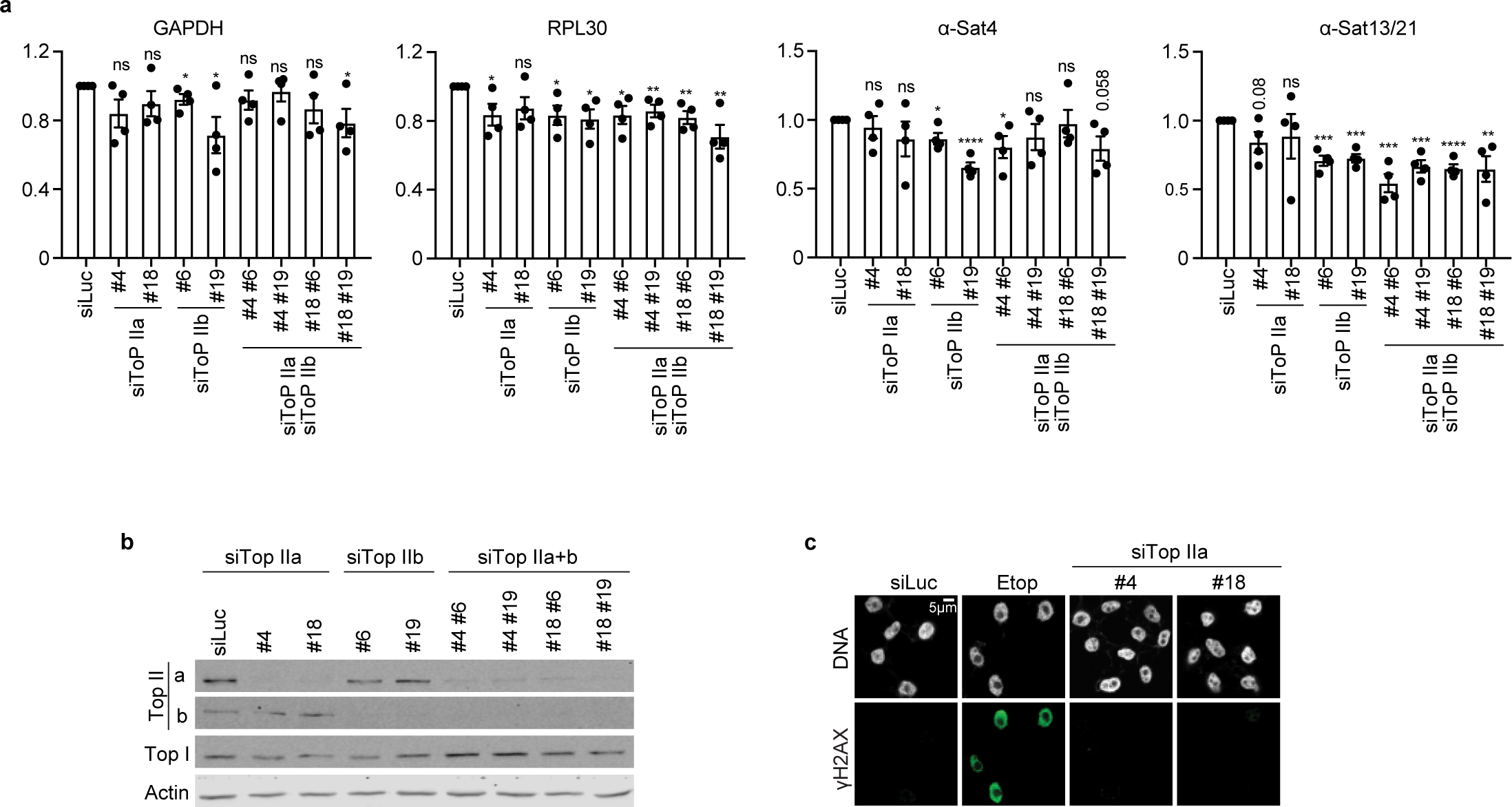
TopII knockdown by siRNAs does not affect α-satellite transcription. **A. TopII knockdown barely changes α-satellite transcription.** RNAs from HeLa cells transfected with the indicated siRNAs were subjected to real-time PCR analysis. **B. Protein levels of TopII in TopII knockdown cells.** Lysates of HeLa cells in (**A**) were subjected to Western-blotting analysis. **C. TopII knockdown does not significantly induce DNA damage**. HeLa cells were transfected with the indicated siRNAs and then subjected to immunostaining with the indicated antibodies.

**Figure S8.**
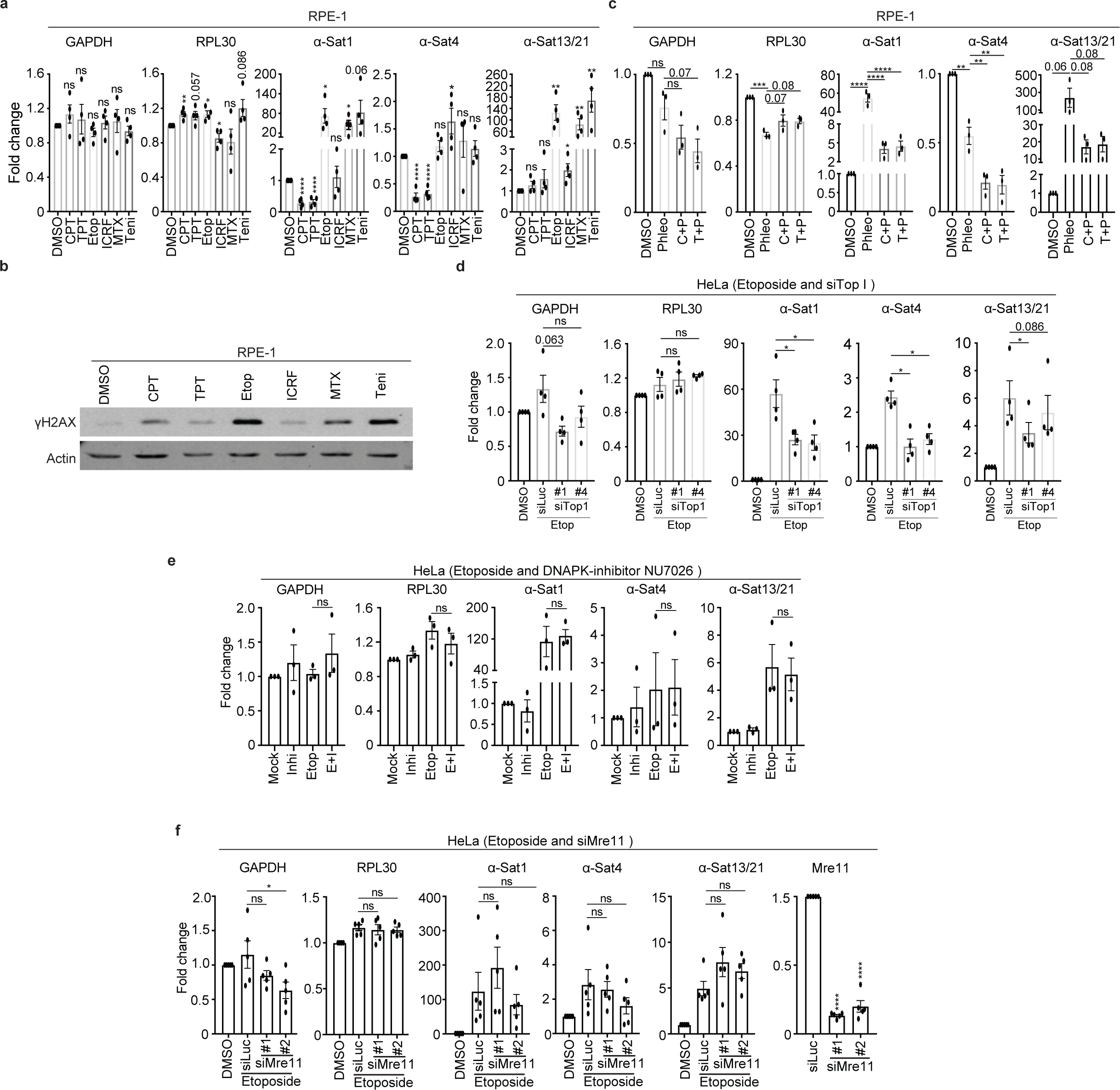
TopI regulates DSB-increased α-satellite transcription. **A. DSB increases α-satellite transcription in RPE-1 cells.** RNAs from RPE-1 cells treated with CPT (2.5 μM), TPT (10 μM), etoposide (Etop, 30 μM), ICRF (20 μM), methotrexate (MTX, 2.0 μM), or teniposide (Teni, 2.5 μM) for 12 hrs were subjected to real-time PCR analysis. **B. γH2AX levels in (A).** Lysates of RPE-1 cells in (**A**) were subjected to SDS-PAGE resolution and then blotted with the indicated antibodies. **C. DSB-induced α-satellite transcription depends on TopI.** RNAs from RPE-1 cells treated with DMSO, phleomycin (Phleo) (24 hrs, 80 μg/ml), CPT (12 hrs, 2.5 μM) plus phleomycin (C+P), or TPT (12 hrs, 10 μM) plus phleomycin (T+P) were subjected to real-time PCR analysis. **C. TopI knockdown by siRNAs partially decreases α-satellite transcription.** HeLa cells transfected with luciferase or TopI siRNAs were treated with etoposide (Etop, 30 μM) for 12 hrs and RNAs were extracted for real-time PCR analysis. **E. DNAPK is dispensable for DSB-induced α-satellite transcription.** RNAs extracted from HeLa cells treated with DMSO, DNA-PK inhibitor NU7206 (0.8 μM), etoposide (Etop, 30 μM), etoposide plus NU7206 (E+I) were subjected for real-time PCR analysis. **F. Mre11 is dispensable for DSB-induced α-satellite transcription by etoposide.** HeLa cells transfected with luciferase or Mre11 (#1 or #2) siRNAs were treated with etoposide (30 μM) for 12 hrs. RNAs were extracted for real-time PCR analysis.

**Figure S9.**
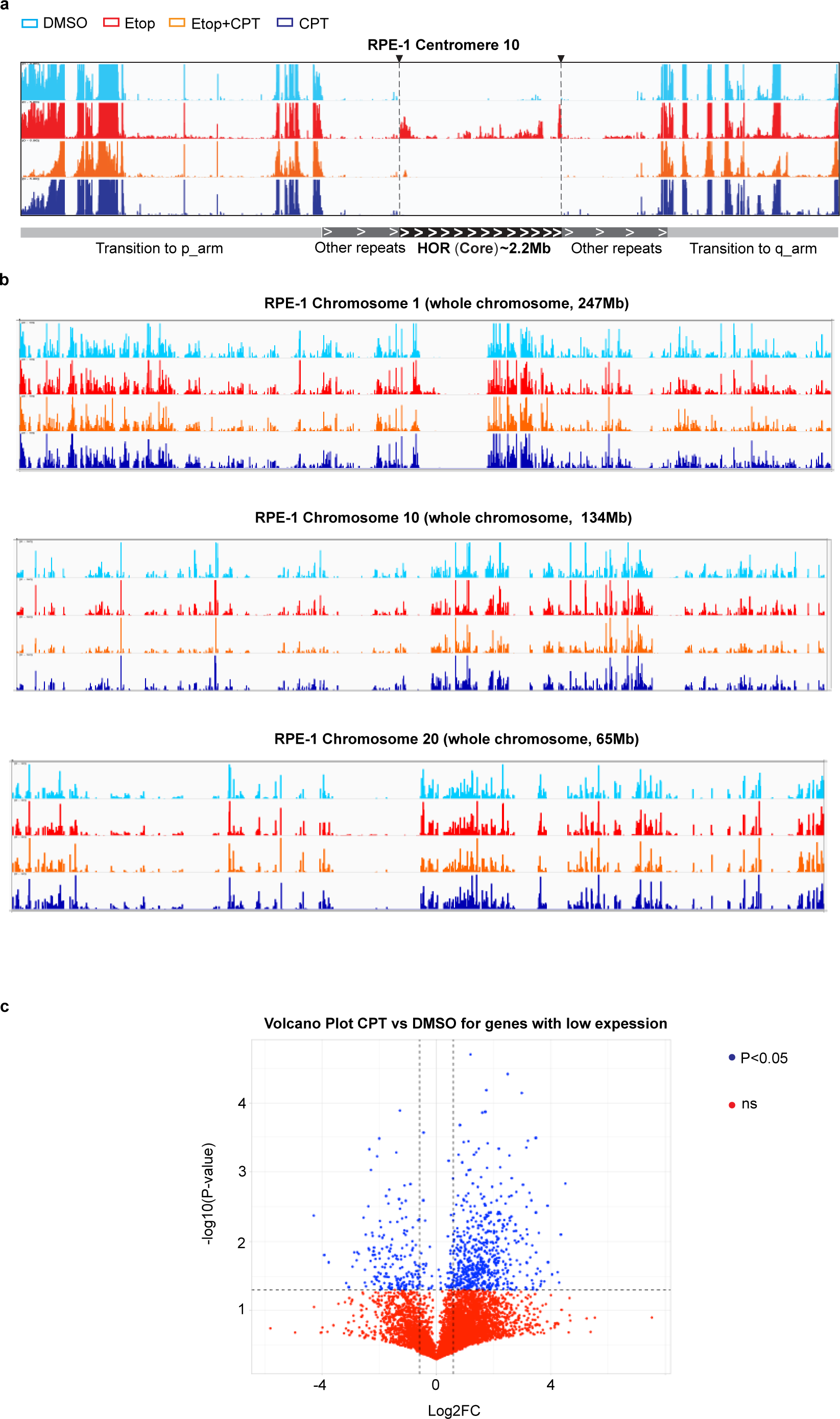
RNA-seq analyses of HeLa cells with treatments of various chemicals. RPE-1 cells were treated with DMSO, CPT (2.5 μM), Etoposide (Etop, 30 μM), or etoposide plus CPT for 12 hrs and total RNAs were extracted for RNA-Seq analysis. Technical details were recorded in the section of Methods and Materials. **a. RNA-seq analysis of α-SatRNAs for centromere 10.** Results for centromeres 1 and 20 were recorded fig.3e. **b. CPT treatment does not significantly change gene expression profile.** RNA-seq analyses were performed for gene expression profile on chromosomes 1, 10 and 20. **c. Analysis of changes in the expression of lowly transcribed genes between CPT and DMSO treatments.** RNA-seq analyses were performed to examine the transcriptional change for lowly transcribed genes after CPT treatment. Details were recorded in the section of Methods and Materials.

**Figure S10.**
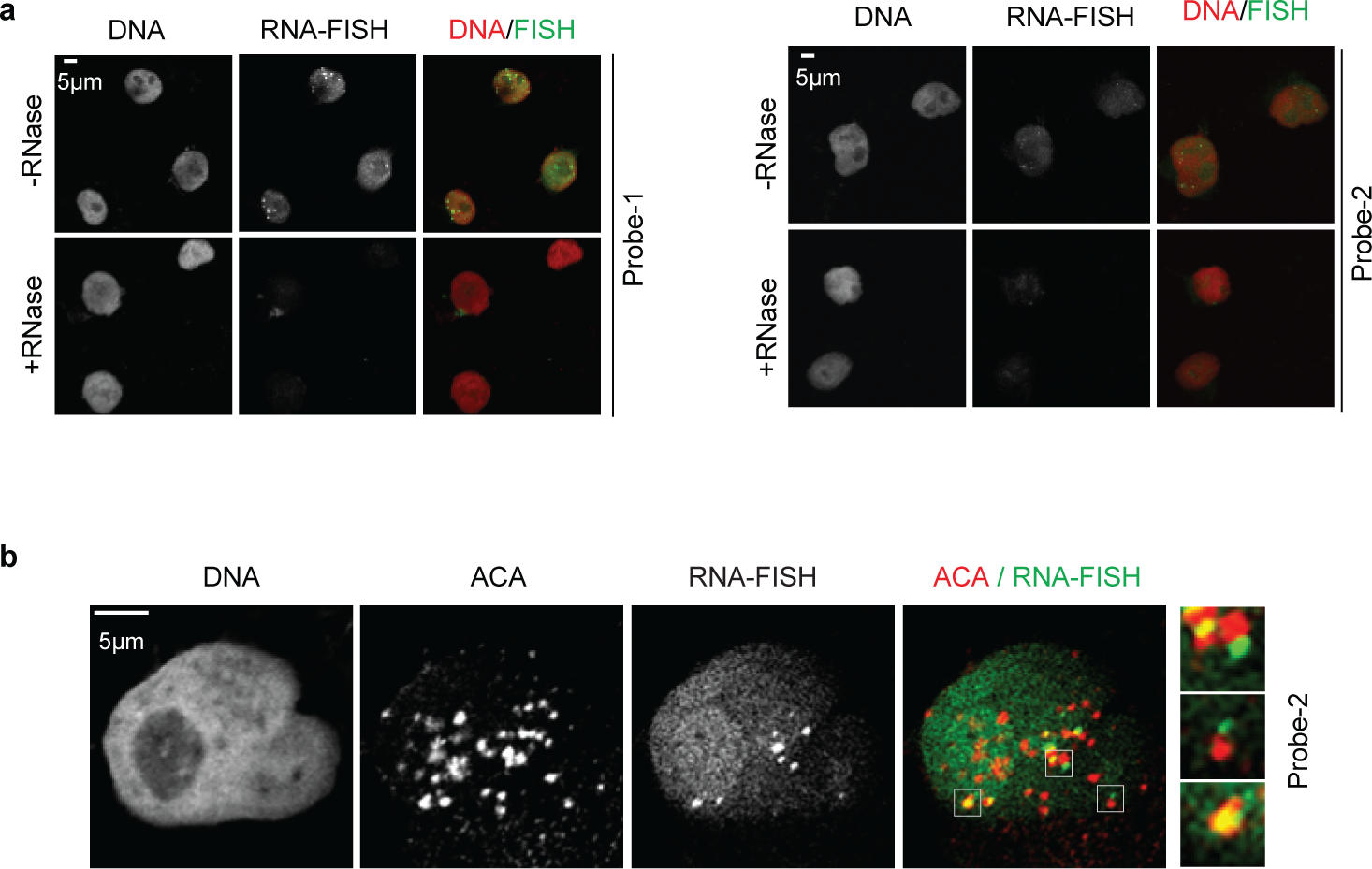
α-satRNA-FISH analyses of HeLa cells with various treatments. **A. α-satRNA-FISH fluorescence signals are sensitive to RNase treatment.** Etoposide-treated (30 μM, 12 hrs) HeLa cells were incubated with RNase before being subjected to hybridization with fluorescence-labelled probe-1 (upper) and probe-2 (bottom). **B. DSB-induced α-satRNA-FISH foci do not always localize to centromeres.** Etoposide-treated (30 μM, 12 hrs) HeLa cells were hybridized with probe-2 and stained with ACA. Selected regions were amplified and placed in the right panel.

